# Longitudinal Habituation and Novelty Detection neural responses from infancy to early childhood in The Gambia and UK

**DOI:** 10.1101/2025.05.23.655801

**Authors:** Anna Blasi Ribera, Borja Blanco Maniega, Samantha McCann, Ebrima Mbye, Ebou Touray, Maria Rozhko, Bosiljka Milosavljevic, Laura Katus, Mariama Saidykhan, Muhammed Ceesay, Tijan Fadera, Giulia Ghillia, Marta Perapoch Amado, Maria M. Crespo-Llado, Sophie E.moore, Clare E. Elwell, Sarah Lloyd-Fox, The BRIGHT Project team

**Author notes:** **Corresponding Author:** Anna Blasi Ribera.

## Abstract

As infants and young children learn from and respond to their environment, their development is driven by their ability to filter out irrelevant stimuli and respond to salient stimuli. While sources and types of stimuli vary across cultural contexts, research to understand the neural mechanisms of these behaviors have largely focused on relatively homogeneous populations in high income settings. To address this lack of diverse representation the Brain Imaging for Global health project (BRIGHT) collected longitudinal data in The Gambia (N=204) and the UK (N=61). Here we present results of the Habituation and Novelty Detection (HaND) fNIRS neuroimaging task. Gambian infants showed persistent response suppression (Habituation) at all visits (from 5mo to 60mo) while Novelty Detection was only observed once infants reached 18 and 24mo. In the UK, infants only showed persistent habituation from 5–12mo, while the response was not evident at 18 and 24mo. Furthermore, in contrast to The Gambia, alongside the habituation patterns observed Uk infants showed novelty detection from 5-12mo. This is the first longitudinal description of the HaND response in individuals from different contextual backgrounds across such a broad age range and number of time points, revealing different patterns of specialization in The Gambia and UK.

**Highlights:**

- This is the first characterization of Habituation and Novelty Detection (HaND) responses in infants and young children from contrasting settings across low and high-income countries.
- In the Gambian cohort, Habituation was typically observed from 5 to 60months; only weak patterns of Novelty Detection were observed at 18 and 24months.
- In the UK cohort, HaND responses were observed up to 12months of age; but were not evident at 18 and 24months.
- Developmental trajectories of HaND responses follow different patterns of specialization from birth to five years of age in individuals living in different contexts.

## 1. Introduction

Interaction with our immediate context is essential for development from a very young age. Infants and young children continuously learn from and respond to their surroundings and this development is guided by their ability to filter out irrelevant information and respond to salient stimuli. This bias to prioritise stimuli not previously encountered (i.e. novelty detection) and suppress responses (i.e. habituation) so as not to expend energy on recurrent information are two core processes of neurodevelopment (Eisenstein, and Smith 2001). While research to understand the neural underpinnings of these behaviors has been well documented across the lifespan (for review see Nordt et al., 2016), research specifically targeting early development (0 – 5 years) has predominantly focused on a narrow age range, often cross-sectional, and with relatively homogeneous populations in high income settings.

*Habituation*, is a marker of information processing in the central nervous system and of behavioural plasticity; as such, it is tightly linked with learning and is evident across a range of species from sea slugs (Castellucci et al. 1970) to primates (e.g., Baylis and Rolls 1987, Earl K.Miller, Paul M. Gochin, andcharles G. Gross 1991). It has been proposed as a predictor of cognitive development and outcomes in term and pre-term infants (Sicard-Cras et al. 2022). *Novelty Detection* refers to the recovery of the neural response with the presentation of a salient, new stimulus. From birth, human infants have been shown to behaviourally habituate to repeating stimuli and detect a change in stimuli (Carolyn Rovee-Collier and Kimberly Cuevas 2008), such that this is a common approach to employ in developmental science to understand pre-verbal infants’ ability to discriminate changes in stimuli that they touch, smell, see, hear and taste.

Repetition suppression paradigms using a habituation and novelty detection (HaND) approach can also be used to better understand neurodevelopmental processes in perception and cognition, especially in pre-verbal infants (Nicholas B.Turk-Browne, Brian J. Scholl, and Marvin M.chun 2008). This neuroscientific approach has two major advantages (Nordt, Hoehl, and Weigelt 2016), firstly the same paradigm can be applied across a range of ages - thus it is ideally suited to explore trajectories of neural specialization across early development – and second, it can be conducted without requiring an overt behavioural or verbal response and so can be applied across a range of contexts (i.e. preterm infants, clinical and cross-cultural comparisons). HaND-based paradigms have been applied using a range of imaging methods to examine cognitive development (Sicard-Cras et al. 2022; Cortesa, Hudac, andmolfese 2019). The characteristic pattern of brain response to the HaND paradigm was first described in 3 to 4month-old sleeping infants with fNIRS as a reduction in neural signal amplitude with stimulus repetition to auditory tones, followed by recovery upon stimulus change (Nakano et al. 2009). Later, Bouchon, Nazzi and Gervain (2015) demonstrated that modifying the complexity of auditory stimuli altered newborn responses to a HaND paradigm on the left hemisphere indicating left hemisphere specialization to linguistic regularities from birth (Bouchon, Nazzi, and Gervain 2015). In the context of learning and recollection from memory, Benavides-Varela et al. (2011) applied the HaND paradigm to explore the role of music and words in memory formation, demonstrating that newborns recognized previously heard words after silence or music, but not after unrelated words (Benavides-Varela et al. 2011).

Interestingly, Dehaene-Lambertz, Hertz Pannier et al. (2006) found that if using a more complex sentence, with only one repetition, in an fMRI paradigm 3-month-olds displayed repetition enhancement in contrast to adults who showed neural repetition suppression (Dehaene-Lambertz et al. 2006). They suggest that the one repetition combined with the complexity of the stimuli may have led to infants still being in a phase of learning and therefore revealing enhanced neural activation. Therefore, the change in, and complexity of, the stimuli may have ramifications for the neural responses observed and may differ across ages. As infants begin processing the complexity of their environment from birth, their responses to auditory stimuli adapt through learning and experience, which influence how efficiently they represent their surroundings (Willmore and King 2023). By studying context-dependent brain responses to auditory stimuli at birth we can begin to understand how these may potentially shape developmental trajectories of the response to novelty as postulated by Háden et al. (2013) in a study with Event Related Potentials (ERPs). The literature on the neurodevelopmental trajectory of the auditory HaND response across early childhood however is limited. We found only one study that used ERPs with an oddball paradigm to examine changes in the response from infancy over time. This longitudinal study included visits at 6months, 2 years, and 4 years of age involving a cohort of 43 children from the Montreal area in Canada (Deguire et al. 2022).

In addition to changes in stimuli, other environmental contextual factors that may have an influence on the neurodevelopmental trajectory of the HaND response include child level factors (i.e. prematurity) and family or community level factors (i.e. poverty associated factors such as undernutrition or culturally influenced family caregiving factors). Bisiacchi, et al (2009) found that neural auditory processing using an oddball paradigm in premature newborns was impacted if the infants were born prior to 30 weeks gestation, as revealed by a lower ERP response, relative to those premature newborns born after 30 weeks. Family or community level factors that influence this neural auditory response have been relatively unexplored during early development, particularly outside of high-income settings. Given the diversity of cultural contexts and environmental stimuli worldwide, findings from such research are often not generalizable. Consequently, factors that drive this specialization remain underrepresented (Basnight-Brown, Janssen, and Thomas 2023). In previous work we have observed that the auditory oddball ERP paradigm showed differences in habitation efficiency in a Gambian cohort of infants relative to a UK cohort (Katus et al. 2020). However, this longitudinal assessment was only conducted at 1, 5 and 18months of age and so we do not know how these findings map onto different ages in early development or different stimulus complexity. Consequently, it is imperative to expand this field further, to underrepresented populations in under-resourced contexts. To the best of our knowledge, no studies have yet explored the neurodevelopmental trajectory of the HaND task; across a large number of age points with a consistent stimulus paradigm to fully understand early developmental changes from 0 – 5 years and in children living outside of a high resource context within a high-income country.

Within the last decade, functional Near Infrared Spectroscopy (fNIRS) has increasingly been employed in neurodevelopmental research beyond high income settings (Blasi et al. 2019; Ayaz et al. 2022; Fishell 2020). The main motivations behind this shift are twofold: (1) the need to implement fNIRS and other imaging techniques in low- and middle-income countries (LMICs) to gain objective insights into neural mechanisms involved in development beyond anthropometric and behavioural assessments; and (2) to address the evident underrepresentation of participants from under-resourced communities in developmental science (Basnight-Brown, Janssen, and Thomas 2023; Henrich, Heine, and Norenzayan 2010). For this, fNIRS systems have been successfully deployed to a wide range of settings to investigate how local contextual factors influence cognitive development (Lloyd-Fox et al. 2015; Jasińska and Guei 2018). fNIRS paradigms originally validated in high income countries have been adapted for cultural relevance and successfully implemented in LMICs (Blasi et al. 2019; Katus et al. 2019). Studies in The Gambia (Sarah Lloyd-Fox et al. 2015; S. Lloyd-Fox et al. 2017), Bangladesh (Perdue et al. 2019; Pirazzoli et al. 2022), and in rural India (Wijeakumar et al. 2019) demonstrate the feasibility of using fNIRS to assess cognitive function in young infants. These studies demonstrate (1) the versatility of fNIRS technology; (2) its capacity to measure associations between psychosocial parameters and neural activity; and (3) that much remains unknown about these mechanisms.

The Brain Imaging for Global Health project (BRIGHT, globalfnirs.org/the-bright-project) provides a platform to examine these responses across different cultural contexts using the HaND paradigm. BRIGHT is a longitudinal, two-site study trackingchildren from gestation to 2 years of age in The Gambia (GM) and the United Kingdom (UK), with an additional pre-school assessment in The Gambia at 3-5 years of age (Sarah Lloyd-Fox et al. 2024). The project was designed to map developmental trajectories of cognition and brain function in contrasting settings. Rather than making direct comparisons between countries, the focus was on intra-country developmental patterns. Also, given that some of the paradigms were novel in all contexts, a UK cohort was essential for methodological validation (Sarah Lloyd-Fox et al. 2024).

Socio-cultural differences between The Gambia and the UK - such as household size, exposure environments, and family interactions (see point 2.1 of the methods section for a description of the GM and UK populations) - may have influenced how infants responded to the stimuli. Therefore, the stark contrast between the UK and Gambian socio-economic contexts makes it highly likely that distinct developmental trajectories could emerge. Preliminary results from the fNIRS (HaND) task at 5 and 8months (Lloyd-Fox et al. 2019) and earlier EEG data (Katus et al. 2020) revealed differing developmental trajectories: Gambian infants exhibited habituation but no evidence of novelty detection, while UK infants showed robust novelty detection after a period of habituation. Here we extend these findings to include data up to 24months in the UK, and up to pre-school age in The Gambia.

Our objectives are threefold: (1) to describe the developmental trajectory of HaND responses in both cohorts from 0 – 2 years; (2) to characterize responses in the Gambian cohort through early childhood (to 5 years) to better understand how contextual differences can shape developmental trajectories; and (3) to capture spatial and temporal brain activity with higher resolution. To ensure consistency, we initially used the same region of interest (ROI)-focused analysis from our previous work, based on a simplified Cluster Permutation Analysis (CPA). Additionally, we employed Threshold-Free Cluster Enhancement (TFCE) analysis to evaluate all available channels and full hemodynamic responses, without requiring pre-defined time windows.

We hypothesized that (1) habituation to repetition of stimuli would occur in both cohorts, (2) novelty detection would emerge in the Gambian cohort similarly to the UK cohort, although at later developmental stages (given our preliminary earlier findings with 5 - 8month olds – Lloyd-Fox et al. 2019); and (3) similarly to the ERP study by Deguire et al. (2022) the responses may become weaker at later ages due to lack of interest or attention to the task or due to rapid repetition suppression (as shown in adults neural responses to repeating sentences; Dehaene-Lambertz et al. 2006). This study primarily describes the observed neural responses in each population, while further investigations into the factors driving these differences within each population will be addressed in future work. Understanding the HaND response in the Gambian sample will help theoretically understand how contextual differences can shape developmental trajectories and will provide practical information by identifying risk and resilience factors across both contexts so we can start to consider interventions that could protect at risk infants.

## 2. Methods

### 2.1. Participants

A total of 204 infants in The Gambia (GM, cohort 1) and 61 infants in the UK (cohort 2) were enrolled in the BRIGHT study at 1month of age onwards (see (Sarah Lloyd-Fox et al. 2024) for a full description). Families were enrolled during pregnancy, and the infants were seen at follow up visits at 1, 5, 8, 12, 18 and 24 months of age. All participating infants were awake during the fNIRS session, except for the 1-month age point. To avoid potential confounding effects of sleep on the haemodynamic response to the stimulation, here we present results at 5months and above only. The infant’s age at each visit ranged from plus or minus two weeks from the targeted age; hereafter we will identify each visit by its targeted age (i.e. visit 1 will be *5mo*). At both sites, testing started in June 2016 and ended in May 2019 in the UK and in July 2020 in The Gambia. Gambian infants were further tested in a single visit at 3 to 5 years of age between August 2021 and March 2022 (as part of the BRIGHT Kids extension of the initial study).

The Gambia is situated on the Atlantic coast in West Africa and is the smallest country in continental Africa. The population of The Gambia is relatively young, with 45% aged 0-14 years, and only 4% aged 65 or older. Approximately half of the population have no formal education, and 56% fall into the lowest wealth bracket. A significant proportion of women (68%) work in agriculture (Gambia Bureau of Statistics (GBoS) and ICF 2021). The BRIGHT participants were from the rural region of West Kiang, where households are typically multi-generational and large, averaging 16 people per compound (Hennig et al. 2017). In contrast, the UK participants lived in and around the city of Cambridge. Most residents in this region live in urban areas, with an average household size of 2.3 people. Education attainment is high: 36.7% of people 16 and above hold post-graduate degree or equivalent, while only 16.7% have no formal qualifications. The rate of unemployment is low, at just 2.3% (https://www.plumplot.co.uk/Cambridgeshire-census-2021.html).

The BRIGHT study was conducted according to guidelines of the Declaration of Helsinki: informed consent was obtained from the families before starting the sessions and ethics approval were obtained from the corresponding committees before recruiting started. Cohort 1 families were recruited from the rural Kiang West District in The Gambia, the majority from the village of Keneba and neighboring villages. Families were identified using the Kiang West Demographic Surveillance System (Hennig et al. 2017; Gambia Bureau of Statistics (GBoS) and ICF. 2021. 2021). The BRIGHT study was approved by the joint Gambia Government-MRCG Ethics committee (ref # SCC 1451), and its follow up BRIGHT Kids was approved by The Gambia Government/MRC Joint with MRC Gambia at LSHTM (Project reference 22737); informed consent was obtained from carers/parents in writing or via thumbprint if they were unable to write. Cohort 2 families with healthy pregnancies were given information about the study at one of their 32 to 36-week gestation ante-natal appointments at the Rosie Hospital, Cambridge University Hospitals NHS Foundation Trust. Those who had expressed interest were contacted via phone call or email and recruited into the study. Families lived in rural or urban communities within a 20-mile radius surrounding the university town of Cambridge. The study was approved by the National Research Ethics Service Committee East of England (REC reference 13/EE/02000), and informed written consent was obtained from parents of infants prior to participate.

Inclusion criteria to the study for both cohorts required that the infants were born at term (37 to 42 weeks gestation). In the UK, a minimum birth weight of 2.5 kg was also required for inclusion and participating caregivers needed to understand English to participate. Ethnicity in the West Kiang District of The Gambia is predominantly Mandinka (79.9% of the population (Hennig et al. 2017)), with its unique language and cultural characteristics. To avoid confounds caused by multiple translations of the study protocols, GM families recruited into the study were only of the Mandinka ethnicity.

Anthropometric measures (weight, length, head circumference) were collected from both cohorts using standardized protocols and equipment and were then converted to z-scores using WHO child growth standards: head circumference (HCZ); weight-for-height (WHZ); and body mass index-for-age (BMIZ): methods and development (2006, ISBN: 924154693X). These measures were used here to exclude infants with severe growth faltering: infants with more than two or more consecutive visits with a score <-3 standard deviations (SD) of the WHO Growth Standards median would be excluded from the analysis. Figure 1 summarizes the number of participants included at each visit.

**Figure 1.**
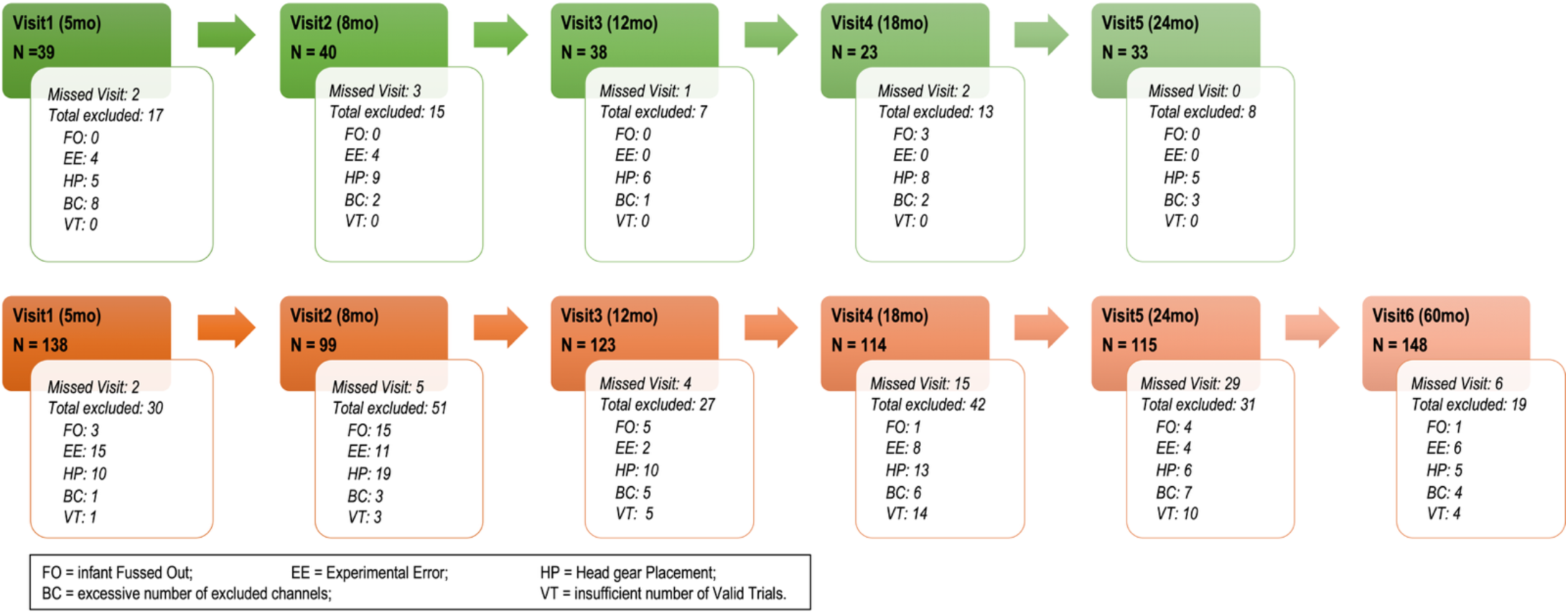
Number of datasets included in the analysis at each time point (N), per cohort. Also, infants missing each visit and the total number of excluded datasets, including the reasons for exclusion, are specified for each time point and cohort.

### 2.2. Experimental Procedures

fNIRS data were collected with the NTS optical topography system (Gowerlabs Ltd., UK, source wavelengths of 850 and 780nm) while participants sat on their carer’s lap. Participants wore a custom-made headgear consisting of two 17-channel arrays configured with 12 sources and 14 detectors (Figures 2 and 3). Before starting the session, head measurements were taken (head circumference, ear-to- ear measurements over forehead and over the top of the head) to guide alignment of the headgear with the 10/20 system anatomical landmarks (Sarah Lloyd-Fox, Richards, et al. 2014), and were used as inputs to the co-registration algorithms that helped calculate the sensitivity profiles of the fNIRS arrays (Collins-Jones et al. 2021). With the headgear securely placed on the participant’s head and before the start of the recording, photographs were taken of the infants wearing the headgear. Whenever possible, photographs were taken at the end as well. The pictures were used in the assessment of headgear placement (Blasi et al. 2014). Measurements on the pictures were also part of the co-registration process, which, with the help of age-appropriate brain templates, identified the brain regions of sensitivity for each channel. These ranged from the inferior frontal gyrus at the front of the head, to the posterior sections of the superior, middle, and inferior temporal gyri (for more details, see Table S1 of the Supplementary Information). To ensure consistent positioning of the headgear across participants, the same protocol for headgear placing was identical at both sites and used the same anatomical landmarks. Prior to data analysis, headgear position was assessed from pictures by the same team of researchers. Due to warping of the pictures, this assessment involved measuring displacement of a reference optode (the middle optode in the lower row on either side of the head) along horizontal (x) and vertical (y) axes, overlaid on each photograph. The x-axis was aligned with the anterior-posterior line defined by the top of the ear and the highest point in the eyebrow; and the y-axis was defined along the line from the tragus to the upper point where the ear joins the head. Displacement was measured using the known inter-optode distance as scaling factor, and datasets were excluded from further analysis when lateral or vertical displacement of the reference optode exceeded ±1.6 cm (Blasi et al. 2014). At 60mo (GM) the custom-made band holding the optode arrays was replaced by a modified Easycap (©GmbH) which facilitated replicability of headgear positioning with the older participants. At all visits, datasets with no pictures available for headgear placement measurements were discarded from the analysis.

**Figure 2.**
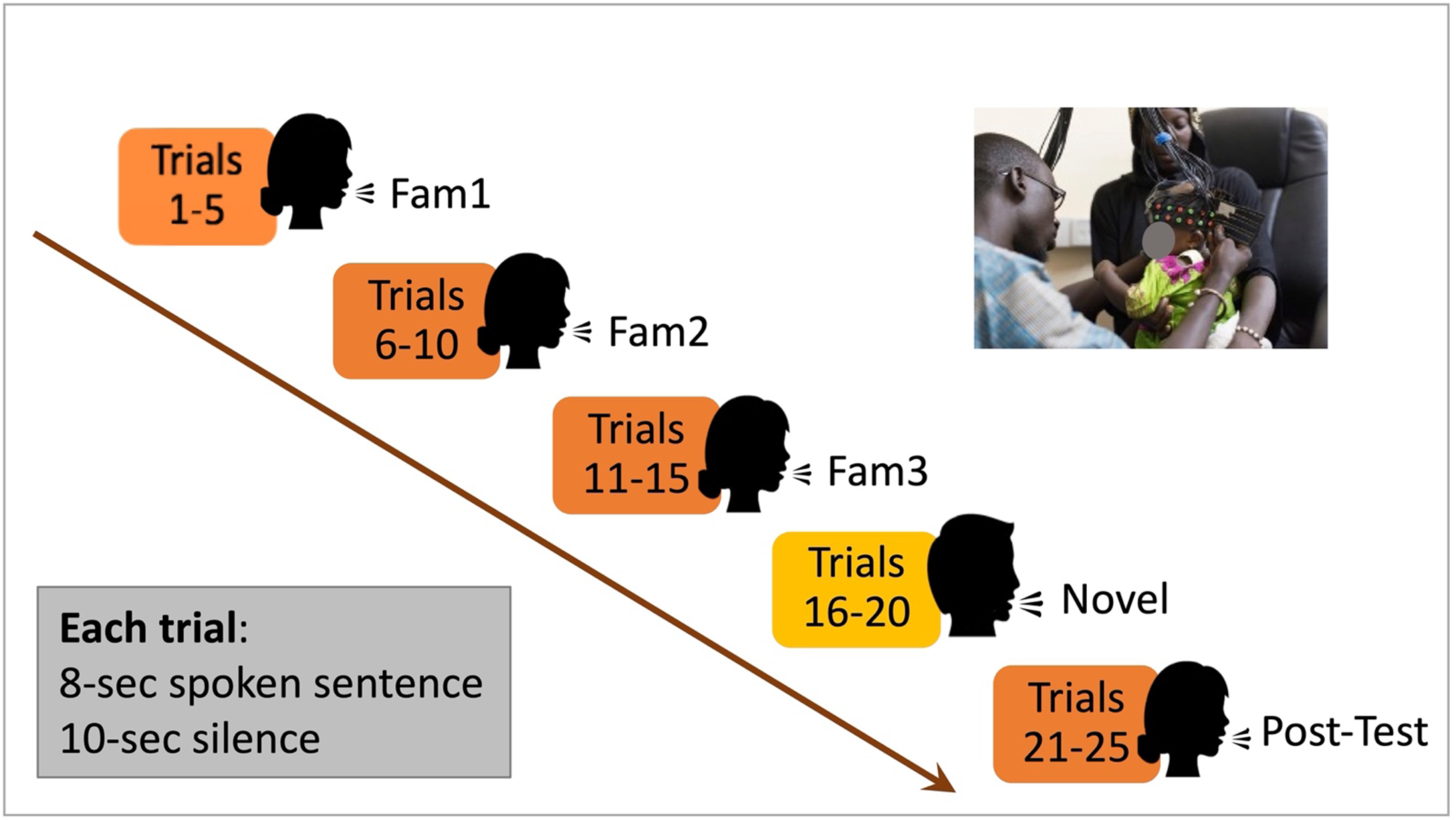
Experimental paradigm, showing one participant being fitted the fNIRS headgear, the sequence of stimuli presentation and the stimulation and baseline timings. Picture: Ian Farrell ©.

**Figure 3.**
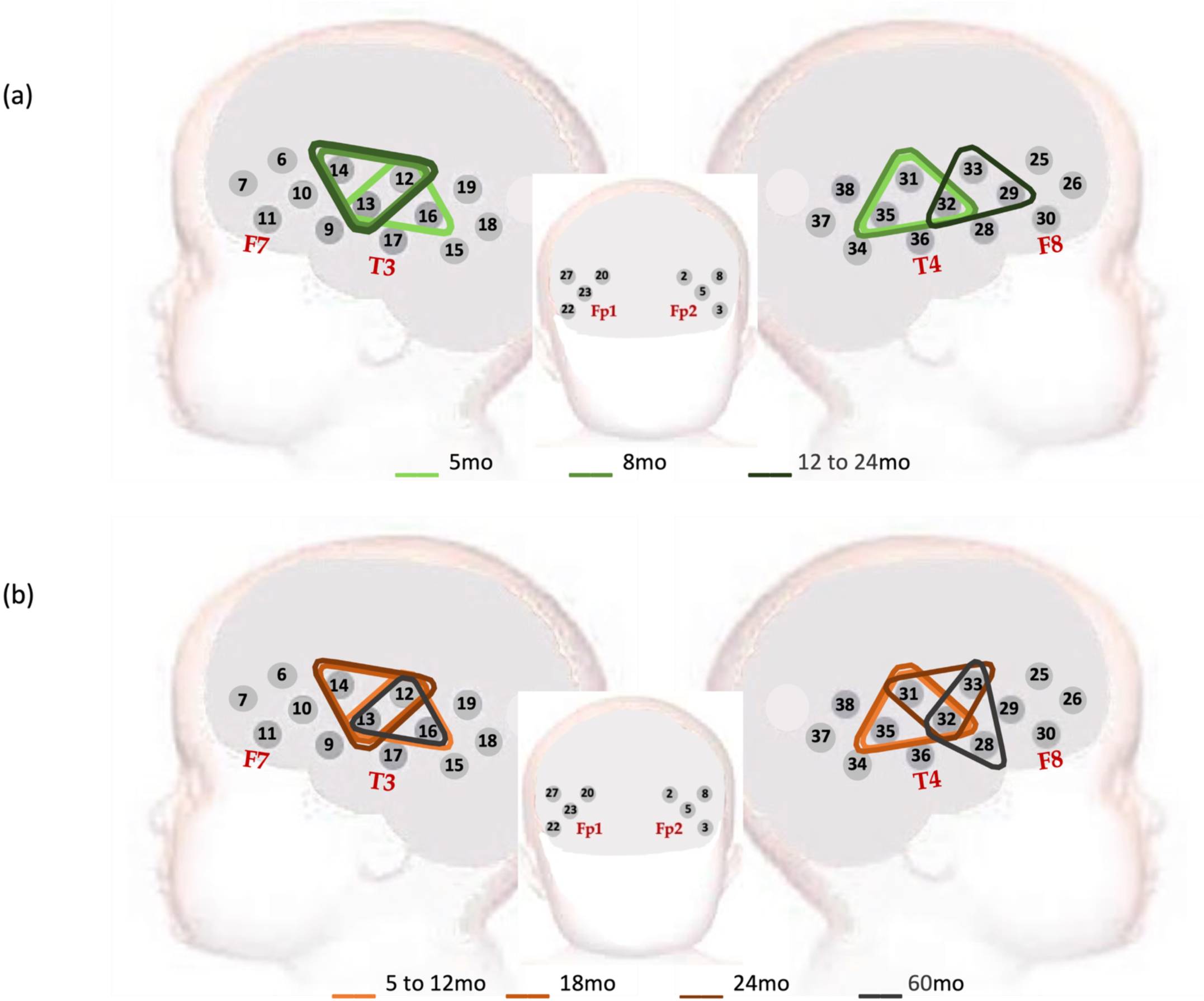
Representation of the channel positions on the participants’ head. The source optodes for channels 17 (on the left hemisphere) and 36 (on the right), in anterior positions to the corresponding channel, were used as references for positioning the head gear before starting the session. These optodes were also used to assess headgear placement on each participant. The triangles represent regions of interest (ROI) calculated with cluster permutation analysis for cohort 1 (in shades of green) and cohort 2 (in shades of orange), for the Fam1 epoch. Merged age groups indicate a shared ROI resulting from CPA analysis.

Carers were encouraged to refrain from interacting with the infant during the data acquisition. The fNIRS session consisted of multiple tasks run consecutively whenever possible: first, the Social task (Sarah Lloyd-Fox, Papademetriou, et al. 2014) with a duration of about 6min; followed by 4min of Functional Connectivity data acquisition (Bulgarelli et al. 2024); after that, the HaND task was run with a total duration of 7min and 30sec; finally the functional connectivity section was repeated. The session was paused if the participant became too fidgety or upset. Within the HaND task, any dataset with a pause within the first 15 trials lasting longer than a few minutes was excluded from the analysis (these infants were classed as a “Fussing out” (FO), see Figure 1). To keep the infants in a stable attentional state, relaxed and focused during the task, an experimenter blew bubbles in front of them without directly engaging in any social interaction. At older ages (from 18mo of age), the bubbles no longer served to engage the infants’ attention and were switched to soft books or silent toys.

The experimental paradigm consisted of repeated presentations of a spoken sentence. These recordings were made of colleagues of the BRIGHT research team who were not involved in the study. They were asked to speak as if they were speaking to a baby. In the UK, the English version of the sentence was used: “*Hi baby! How are you? Are you having fun? Thank you for coming to see us today. We’re very happy to see you*”. In The Gambia, this was translated to Mandinka as: “*Denano a be nyadii. I be kongtan-rin? Abaraka bake ela naa kanan njibee bee, n kontanta bake le ke jeh*”. For more details on the recorded voice format and the playback set up see (Lloyd-Fox et al. 2019). The sentence, with a duration of 8sec, was repeated 25 times, with a silent baseline of 10sec between two consecutive presentations. The session started with at least 10sec of silent baseline. A female voice spoke the first fifteen presentations of the sentence (trials 1-15); then it was switched to a male voice for five repetitions (trials 16-20); the initial female voice returned for the final five presentations (trials 21-25). One trial was defined from condition onset to the end of the following baseline. The trials were grouped in epochs, each with 5 consecutive trials: Familiarization epochs included trials 1-5 (Fam1), 6-10 (Fam2) and 11-15 (Fam3); Novel epoch included trials 16-20; and Post Test, trials 21-25 (as illustrated in Figure 2).

Data quality was assessed at each channel and those channels that did not meet the criteria (see section 2.3, Data Processing) were discarded from further analysis. In line with all our previous publications, datasets with more than 40% of discarded channels were excluded from further analysis.

The infants’ behavior was recorded during the session and their engagement with the task was coded to discard trials with behaviours that could potentially impact on their responses, such as distraction with their surroundings, interacting with the carer, tiredness or unhappy mood. Epochs with less than 60% infant engagement were excluded from further analysis. Note that to be included in the analysis, datasets were required to have all three familiarization epochs with more than 60% of valid data in each. Data exclusion for insufficient number of Valid Trials (VT) in Figure 1 includes infant behavior as well as trials excluded for excessive motion artefacts (see the data processing section for details). If the video recording was missing, notes about infant behavior taken by the experimenters during the session were consulted to determine the suitability of including the dataset; if no notes were available or if an overall lack of engagement with the study was reported in the notes, the dataset was excluded.

### 2.3. Data processing

Time courses of all channels were visually inspected for potential experimental errors and, at this point, datasets with issues such as missing event markers, or an excessive number of channels with light saturation were excluded. Metadata information such as behavioral coding and headgear placement were compiled separately and added during pre-processing.

The fNIRS datasets were pre-processed with the set of in-house Matlab® scripts used in previous publication (Lloyd-Fox et al. 2019). The pre-processing pipeline included: (1)channel pruning to remove channels with poor signal (intensity<3e-4), using the coefficient of variation (with a maximum threshold of 0.2 for both wavelengths), and a maximum power of 0.045 in the (0.04 to 0.4)Hz frequency band; (2) low-pass filtering with a cut off frequency 0.8Hz to remove high-frequency oscillations including heart rate; (3) segmentation into trials, each one starting at t = -4sec and ending at t = 18sec, with t = 0 being the time of stimulus onset; (4) conversion of the optical density signals into oxy- and deoxy-haemoglobin (HbO2 and HbR, respectively) with the modified Beer-Lambert law (Delpy et al. 1988), using a differential pathlength factor (DPF) that varied with age of the participants (Duncan et al. 1995) to take into account age-related changes in brain tissues; (5) within each trial and channel, motion artefacts detection and trial removal if the HbO2 signal exceeded ±3.5μMolar at baseline prior to the stimulus onset or ±5μMolar during stimulation; (6) baseline detrending by fitting a straight line between the average of the 4sec at the start and the end of each trial; (7) block averaging across valid trials within each epoch for group analysis.

Following pre-processing, datasets were excluded from further analysis if there were an insufficient number of valid channels (BC in Figure 1; <60%, the equivalent to less than 21 valid channels) or insufficient number of valid trials (VT). A minimum number of three valid trials were required for an epoch to be considered valid. A dataset missing any of the familiarization epochs (Fam1, Fam2 or Fam3) was excluded from the analysis. However, datasets with missing Novel or PostTest epochs would still contribute to the analysis.

The number of trials contributing to each epoch was similar in both cohorts (Table 2). An observable trend for all age points and in both cohorts is that the number of trials included in the later epochs (Novel and PostTest) is slightly reduced compared to the familiarization epochs. This reflected increasing tiredness/disengagement as the session progressed and was expected. However, the number of infants contributing data to the Novel and PostTest epochs overall remained high.

**Table 1.** Summary of participant characteristics for the UK and Gambian cohorts (mean ± SEM).

| Characteristics | GM Cohort |  |  |  |  |  | UK Cohort |  |  |  |  |
| --- | --- | --- | --- | --- | --- | --- | --- | --- | --- | --- | --- |
|  | 5mo | 8mo | 12mo | 18mo | 24mo | 60mo | 5mo | 8mo | 12mo | 18mo | 24mo |
| Sex (m/f) | 64/74 | 48/51 | 58/65 | 52/62 | 53/62 | 74/74 | 23/16 | 23/17 | 23/15 | 45/78 | 18/15 |
| Age (days) | 161 ± 1 | 250 ± 2 | 378 ± 2 | 581 ± 5 | 747 ± 2 | 1515 ± 215 | 156 ± 1 | 250 ± 2 | 377 ± 2 | 564 ± 4 | 738 ± 4 |
| Weight (kg) | 6.81 ± 0.07 | 7.51 ± 0.09 | 8.32 ± 0.1 | 9.48 ± 0.11 | 10.65 ± 0.12 | 15.09 ± 1.75 | 7.22 ± 0.15 | 8.5 ± 0.16 | 9.69 ± 0.21 | 11.1 ± 0.29 | 12.26 ± 0.3 |
| Height (cm) | 64.28 ± 0.19 | 68.07 ± 0.25 | 72.48 ± 0.25 | 78.75 ± 0.28 | 83.16 ± 0.32 | 102.43 ± 4.94 | 64.44 ± 0.36 | 69.28 ± 0.47 | 74.62 ± 0.45 | 80.69 ± 0.69 | 85.84 ± 0.71 |
| Head circumference (H-C, cm) | 41.38 ± 0.12 | 43.05 ± 0.14 | 44.3 ± 0.15 | 45.72 ± 0.17 | 46.57 ± 0.16 | 49.21 ± 1.42 | 43.05 ± 0.2 | 45.09 ± 0.19 | 46.59 ± 0.18 | 47.85 ± 0.26 | 48.99 ± 0.26 |
| MUAC (cm) | 13.27 ± 0.1 | 13.41 ± 0.11 | 13.56 ± 0.09 | 13.91 ± 0.11 | 14.39 ± 0.1 | N/A | 14.53 ± 0.21 | 15.33 ± 0.23 | 15.51 ± 0.27 | 15.74 ± 0.29 | 16.19 ± 0.36 |
| Growth and anthropometric scores<sup>a</sup> |  |  |  |  |  |  |  |  |  |  |  |
| Weight-for-age (WAZ) | -0.64 ± 0.09 | -0.92 ± 0.1 | -1.08 ± 0.09 | -1.09 ± 0.1 | -0.97 ± 0.09 | -0.69 ± 0.88 | -0.15 ± 0.18 | 0.04 ± 0.17 | 0.14 ± 0.19 | .19 ± 0.24 | 0.19 ± 0.21 |
| Height-for-age (HAZ) | -0.5 ± 0.08 | -0.77 ± 0.1 | -1.12 ± 0.09 | -1.18 ± 0.1 | -1.18 ± 0.1 | -0.32 ± 0.98 | -0.41 ± 0.16 | -0.35 ± 0.19 | -0.35 ± 0.18 | -0.53 ± 0.24 | -0.32 ± 0.23 |
| H-C-for-age (HCAZ) | -0.64 ± 0.08 | -0.71 ± 0.1 | -0.96 ± 0.1 | -0.85 ± 0.11 | -0.84 ± 0.11 | -0.44 ± 0.92 | 0.69 ± 0.15 | 0.73 ± 0.14 | 0.63 ± 0.13 | 0.62 ± 0.22 | 0.9 ± 0.19 |
| Weight-for-height (WHZ) | -0.36 ± 0.09 | -0.61 ± 0.12 | -0.72 ± 0.1 | -0.71 ± 0.1 | -0.52 ± 0.1 | -0.77 ± 0.88 | 0.2 ± 0.2 | 0.42 ± 0.18 | 0.36 ± 0.19 | 0.6 ± 0.24 | 0.51 ± 0.2 |

NOTE: MUAC=Mid upper arm circumference.
<sup>a</sup>With use of WHO reference curves.

**Table 2.** Number of trials per epoch; averages for each age point and cohort (mean ± SEM).

| GM Cohort |  |  |  |  |  | UK Cohort |  |  |  |  |
| --- | --- | --- | --- | --- | --- | --- | --- | --- | --- | --- |
| 5mo | 8mo | 12mo | 18mo | 24mo | 60mo | 5mo | 8mo | 12mo | 18mo | 24mo |
| 4.96 ± 0.02 | 4.97 ± 0.02 | 4.95 ± 0.03 | 4.73 ± 0.06 | 4.71 ± 0.05 | 4.93 ± 0.03 | 4.97 ± 0.03 | 5.00 ± 0.00 | 4.89 ± 0.08 | 5.00 ± 0.00 | 4.82 ± 0.08 |
| 4.99 ± 0.01 | 4.98 ± 0.02 | 4.97 ± 0.02 | 4.81 ± 0.04 | 4.67 ± 0.06 | 4.86 ± 0.04 | 4.95 ± 0.05 | 5.00 ± 0.00 | 4.95 ± 0.06 | 5.00 ± 0.00 | 5.00 ± 0.00 |
| 4.99 ± 0.01 | 4.96 ± 0.03 | 4.99 ± 0.01 | 4.68 ± 0.07 | 4.56 ± 0.07 | 4.73 ± 0.05 | 5.00 ± 0.00 | 5.00 ± 0.00 | 4.95 ± 0.06 | 5.00 ± 0.00 | 4.97 ± 0.03 |
| 4.90 ± 0.05 | 4.82 ± 0.07 | 4.81 ± 0.06 | 4.41 ± 0.11 | 4.37 ± 0.10 | 4.70 ± 0.06 | 4.95 ± 0.05 | 5.00 ± 0.00 | 4.95 ± 0.05 | 5.00 ± 0.00 | 4.82 ± 0.10 |
| 4.68 ± 0.10 | 4.72 ± 0.10 | 4.63 ± 0.10 | 4.23 ± 0.13 | 4.27 ± 0.11 | 4.64 ± 0.07 | 4.79 ± 0.14 | 4.90 ± 0.10 | 4.84 ± 0.14 | 4.78 ± 0.18 | 4.45 ± 0.22 |

### 2.4 Statistical analysis

Two data driven approaches were used to investigate the responses to the HaND paradigm at each visit and epoch: (1) a simplified version of the Cluster Permutation analysis (CPA); and (2) the threshold-free cluster enhancement (TFCE).

*CPA*: This method involved a cluster-based permutation analysis (Maris and Oostenveld 2007) to calculate regions of interest (ROIs). It provides a solution to the multiple comparisons problem where data is collected simultaneously from multiple measurement points close to each other in space and has been successfully used with fNIRS (Benavides-Varela et al. 2017; Ferry et al. 2016; Abboub, Nazzi, and Gervain 2016). Here we used its simplified version for consistency with the preliminary analysis of the data (Lloyd-Fox et al. 2019). The 34 available channels were grouped into 58 pre-defined clusters defined by 3 nearest-neighboring channels, considering only groups arranged in triangular shapes. The cluster-based permutation analysis (CPA) was run for Fam1 and Novel epochs within each cohort and visit to establish ROIs with the strongest initial response to stimuli and to consider that ROIs might differ between these epochs. CPA was run with two time windows to accommodate for potential shifts in the timing of the responses with age: (1) 8 to 12sec post stimulus onset; and (2) 10 to 14sec post stimulus onset (based on the time window selected in (Lloyd-Fox et al. 2019)). The channel composition of the ROIs did not vary with the time window. The largest combined t-value was used to select the time window: 8 to 12sec for visits at 5 and 8mo for the Gambian cohort; and for visits at 5, 8 and 12mo in the UK; 10 to 14sec for all other visits. Either a significant increase in HbO2 concentration, or a significant decrease in HbR, is commonly accepted as an indicator of cortical activation in infant work, and it is best to report both chromophores whenever possible (Obrig and Villringer 2003). However, the largest amplitude changes and highest signal to noise ratio were in HbO2 (S. Lloyd-Fox, Blasi, and Elwell 2010; Pinti et al. 2018; Aslin, Shukla, and Emberson 2015) and so results focused on this signal for the CPA. While HbR offers greater specificity than HbO2, it also has poorer signal to noise ratio (Tachtsidis and Scholkmann 2016; Pinti et al. 2018), and that may negatively affect the performance of CPA.

Mean HbO2 within the pre-defined time windows was calculated for each participant, epoch andchannel. The average across valid channels in the ROI was used as the dependent variable in the General linear model with repeated measures with *Epoch* as fixed effect (including Fam1, Fam3, Novel and PostTest). Post-hoc pairwise comparisons identified *Habituation* (Fam1>Fam3) and *Novelty Detection* (Novel > Fam3) and *Continued Habituation* (Fam3>Novel). Each cohort was analyzed separately. We chose a linear mixed model (LMM) approach with *Visit* as repeated fixed effect, post-hoc pairwise comparisons and Bonferroni correction for multiple comparisons to investigate potential changes over time (differences across age points within each cohort) of *Habituation* and *Novelty Detection*. For each contrast, we selected the model with smallest Maximum Likelihood, Akaike’s Information Criterion, and number of parameters. LMMs are better suited for longitudinal data analysis than the GLM as they can handle correlated data (such as repeated measurements), unequal variances and missing data (Pusponegoro et al. 2017; Geert Molenberghs and Geert Verbeke 2000; Shek and Ma 2011).

*TFCE*: CPA limited our investigation to the 6-channel clusters with the highest t-statistics (where the strongest activation occurred), which were all situated over the middle and superior temporal regions. However, evidence of more frontal *Habituation* and *Novelty Detection* in the literature (Fló et al. 2019; Benavides-Varela et al. 2011; Nakano et al. 2009; Mao et al. 2021), demonstrated the need to investigate activation in the full extent of our sensor array. Therefore, we re-analyzed the data with the TFCE method (Smith and Nichols 2009)). TFCE has been previously used in EEG (Mensen and Khatami 2013), it is available in fMRI processing packages like SPM, and has recently been introduced to the fNIRS community (Blanco et al. 2023; Carius et al. 2023).

The TFCE analysis’ first step is a raw statistical map (condition vs baseline or condition contrast). The next step is a transformation that enhances areas of contiguity (clusters)more than areas with lower statistical values (areas of background noise) in time and space, therefore involving two transformations: the first one enhances the signal with *temporal* contiguity; and the second one enhances areas of *spatial* contiguity. For this step, we set the parameter e=1 and h=2 based on (Pernet et al. 2015) and (Mensen and Khatami 2013). The resulting TFCE map is not intrinsically thresholded, but this enhancement facilitates discrimination between significant regions and background noise. A second step involves permutation testing for statistical inference: a distribution of maximum TFCE values is calculated by permuting the original statistical map, then applying the TFCE temporal and spatial transformations, and from it, selecting the maximum TFCE value. This process is repeated N times (we chose N = 1000) to build the distribution of maximum TFCE values (building the null hypothesis, H_0_). Then, the original TFCE map can be tested against this distribution and obtain the corresponding p-values, using a p<0.025 threshold (2-sided) for epoch significance and epoch contrasts.

We applied TFCE analysis to HbO2 and HbR signals separately, and defined activation as an increase in HbO2 and/or a decrease in HbR; cases of simultaneous increase or decrease in both chromophores were not considered as activation. The TFCE analysis was performed including all 34 channels in the fNIRS arrays, in a time window starting 3sec post-stimulus-onset to the end of the trial (the end of the baseline following each presentation of the stimulus). Epoch contrasts were considered significant only where activation of the first condition was significant. For example, for Habituation, only channels with significant Fam1 activation were further explored, and in those channels, only temporal segments where significant Fam1>Fam3 overlapped Fam1 activation were considered. This method provided clusters and time windows compatible with activation for each epoch and epoch contrast, showing the evolution of the response to the HaND paradigm across time points within each cohort.

## 3. Results

Participant characteristics for each age point and cohort including sex, average age and anthropometry z-scores calculated using WHO child growth standards are summarized in Table 1. The two cohorts differed slightly in the proportion of males/females: in The Gambia, the range was from 46.1% male (24mo) to 48.5% male (8mo); and in the UK, it was from 54.5% male (24mo) to 60.5% male (12mo). In The Gambian cohort, four infants at 5mo, five at 8mo, six at 12mo, three at 18mo, three at 24mo, and four at 60mo had one or more growth standard scores < -3SD from the median WHO reference. However, this did not occur at two or more consecutive visits for any of them. None of the UK infants presented severe growth faltering, and no infants were excluded from the analysis based on their growth scores on either cohort.

Figure 1 summarizes the number of included datasets at each time point. It also illustrates the number of infants that missed each visit, the number of datasets excluded, and the reasons for exclusion. FO indicates that the infant started the HaND task but became upset and the session was stopped before completing the first 15 trials. Experimental errors (EE) included loss of communication between computers or between the computer and the fNIRS system, loss of data due to a storage error, missing pictures for headgear assessment or missing video for behavioral coding.

In this section we first report the results from the CP analysis, the method used to define regions of interest (ROIs, section 3.1). We then report findings using the TFCE analysis (section 3.2), which allowed us to expand the spatial and temporal scopes of our analysis.

### 3.1 Cluster Permutation Analysis (CPA) with a region of Interest (ROI)

Here we report activation to Fam1, Fam3, Novel and PostTest as well as epoch contrasts Fam1 vs Fam3 and Fam3 vs Novel. Activation was measured as significant change in HbO2 from baseline within an ROI at each age point and cohort. A decrease between familiarisation epochs was interpreted as Habituation, and a decrease from Fam1 or Fam3 to Novel was considered as Continued Habituation, as the response continued with the decreasing trend regardless of the change in stimuli.

CPA for Fam1 (or Novel) provided p- and t-values for the pre-defined 3-channel clusters. Then, those with significant activation (p < 0.05) were listed, and from that list, we selected the cluster with the highest t-value per hemisphere. The 6-channel ROI was the result of combining the two clusters. ROIs calculated for Fam1 and Novel were defined by very similar channel combinations (see the Supplementary Information tables S2 and S3 for channel composition of the ROIs for each cohort).

Channel co-registration (Collins-Jones et al. 2021) allowed us to identify the brain regions mapped by each ROI (full details in Table S1 of the Supplementary Information): channels 12, 31 and 32, included in the ROIs for both cohorts and at all visits, were positioned over the superior temporal gyrus (STG) at all time points, while channels 13, 16 and 35 (in the ROI at most time points in both cohorts) were localized over the middle temporal gyrus (MTG).channel 14 was over the STG at 18mo and the pre/post central gyrus at 24mo.channel 33 was localized over the STG at 24mo. The co-registration work was undertaken prior to the completion of data collection at 60mo; therefore, identification of brain regions was unavailable at that visit. Although brain growth is very rapid in the first year of life (from a 35% of adult brain volume at 2-3 weeks after birth to 80% at 24months), it slows after that (Gilmore, Knickmeyer, and Gao 2018). Therefore, we considered the same channel mappings at 24 and 60mo as we assumed it would not change beyond the fNIRS spatial resolution in this period of time.

#### 3.1.1 Cohort 1 (GM)

Gambian infants showed a significant linear effect of epoch at 8 (p=0.006), 12 (p<0.001), 18 (p<0.001), 24 (p=0.024), and 60mo (p<0.001) (see Table 2). Mauchly’s test of Sphericity was met at most visits (p>0.05), except for 24mo, F(5)=11.126, p=0.049) where the Greenhouse-Geisser method was used instead. Post-hoc pairwise comparisons showed that, overall, there was a decrease in the amplitude of the response from Fam1 at all ages. At the younger ages, this decrease was slow: at 5mo, only Fam1>Novel reached statistical significance; and at 8mo, Fam1>Novel and Fam1>PostTest were significant, but not Fam1>Fam3. From 12mo, we observed significant decrease of the response from Fam1 to Fam3 and PostTest, an indication that the response to the familiarization stimuli never returned to the level of the initial response, as evidenced by the significant Fam1>PostTest contrasts. Interestingly, the decrease the amplitude of the response from Fam3 to Novel appeared to reverse at 18 and 24mo as shown in Figure 4b, although the pairwise contrast Fam3 vs Novel was not significant. Moreover, at 24mo it appeared that the amplitude of the responses to Novel and Fam1 were similar as the contrast Fam1>Novel was not significant, unlike at all other points. Although at 18 and 24mo a significant cubic effect of epoch was found (at 18mo, p=0.026; and at 24mo, p=0.029), this analysis was unable to confirm the presence of Novelty Detection in this cohort.

**Figure 4.**
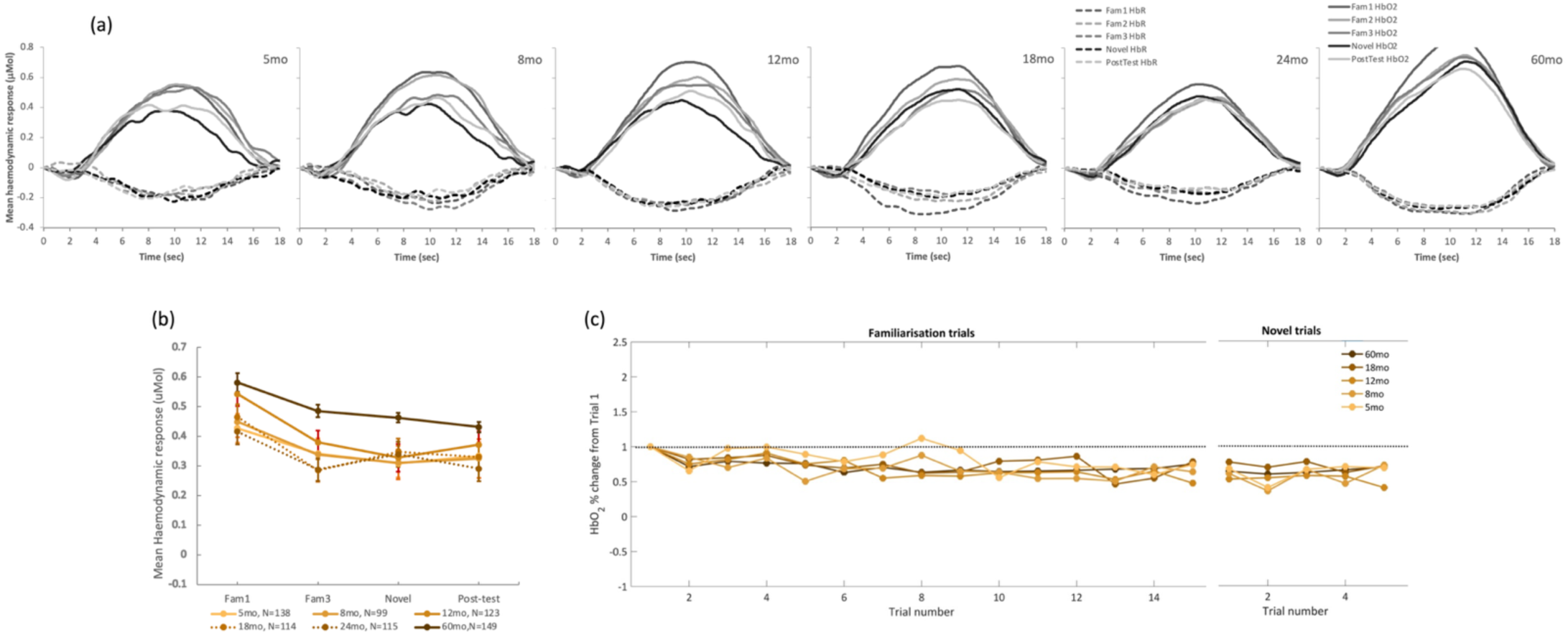
Cohort 1 (GM). Haemodynamic response. (a) Averaged HbO2 (solid lines) and HbR (dashed lines) across all participants at each age point; different epochs are represented by shades of grey. (b) Average HbO2 change from baseline within a time window post stimulus onset (8 to 12sec for 5 and 8mo; 10 to 14 for 12 to 24mo) across valid trials per epoch, at each age point (represented in shades of orange as specified in the legend in Fig3.c). (c) Trial by trial average HbO2 change from baseline within a time window post stimulus onset (8 to 12sec for 5 and 8mo; 10 to 14 for 12 to 24mo), for all familiarization trials (1 to 15) and Novel (16-20) epochs.

To investigate the effect of age on Habituation and Novelty Detection, we applied linear mixed models with visit as repeated effect t. No age effect was observed on either Habituation (F[5,212.847]=1.114; p=0.354) or Novelty Detection (F[5,196.616]=1.469; p=0.201).

Results of the ROI-based analysis for the GM cohort are summarized in Table 3 and figures representing the time course of the responses at all visits, average HbO2 change per trial and per epoch are represented in Figure 4.

**Table 3.** Epoch effect within each visit, GM cohort. General Linear model with repeated measures (Epoch), including post-hoc pairwise comparisons between the most relevant epochs, Bonferroni-corrected for multiple comparisons.

| Visit | N | Epoch effect |  |  | Significant Pairwise comparisons (p-values, Bonferroni-corrected) |  |  |  |
| --- | --- | --- | --- | --- | --- | --- | --- | --- |
|  |  | F | p-val | h2 | Fam1>Fam3 | Fam1 > Novel | Fam1>Post-test | Fam3 > Novel |
| 5mo | 138 | 2.346 | 0.072 | 0.018 |  | 0.057 |  |  |
| 8mo | 99 | 3.429 | 0.018 | 0.037 |  | 0.035 | 0.042 |  |
| 12mo | 123 | 8.077 | <0.001 | 0.067 | 0.003 | <0.001 | 0.005 |  |
| 18mo | 114 | 8.899 | <0.001 | 0.086 | <0.001 | 0.002 | 0.001 |  |
| 24mo | 115 | 3.715 | 0.014 | 0.04 | 0.027 |  | 0.027 |  |
| 60mo | 148 | 15.905 | <0.001 | 0.101 | <0.001 | <0.001 | <0.001 |  |

#### 3.1.2 Cohort 2 (UK)

At 5, 8 and 12mo, infants in the UK showed a response pattern consistent with (1) Habituation from Fam1 to Fam3, and (2) a recovery with the presentation of the Novel stimuli (see Figure 5a and 5b). However, this was not evident at 18 or 24mo. The general linear model analysis with repeated measures using epoch as main factor revealed a significant overall epoch effect at 5mo (F(3)=2.954, p=0.036, η^2^=0.085) and 8mo (F(3)=8.502, p<0.001, η^2^=0.183). At 12mo, epoch did not have a significant linear effect (F(3)=2.024, p=0.115, η^2^=0.056, see Table 4). However, a significant nonlinear (cubic) epoch effect identified the signature of the HaND response at all three visits: 5mo, F(1)=7.964, p=0.008, η^2^= 0.199; 8mo, F(1)=15,229, p<0.001, η^2^= 0.286; and 12mo, F(1)=4.51, p=0.042, η^2^= 0.127 (results summarized in Table 4). At all three visits, sphericity between epochs was met (Mauchly’s Test at 5mo: Χ^2^=0.808, p=0.977; at 8mo: Χ ^2^=3.105, p=0.684; at 12mo: Χ^2^=3.566, p=0.614). Pairwise comparisons between epochs revealed a significantly stronger Novel response relative to Fam3 at 5mo, (p=0.006) and 8mo (p=0.032). Furthermore, at 8mo, response to Fam1 was stronger than Fam3 (p=0.005) and Post-Test (p=0.001). No pairwise significant differences were detected at 12mo.

**Figure 5.**
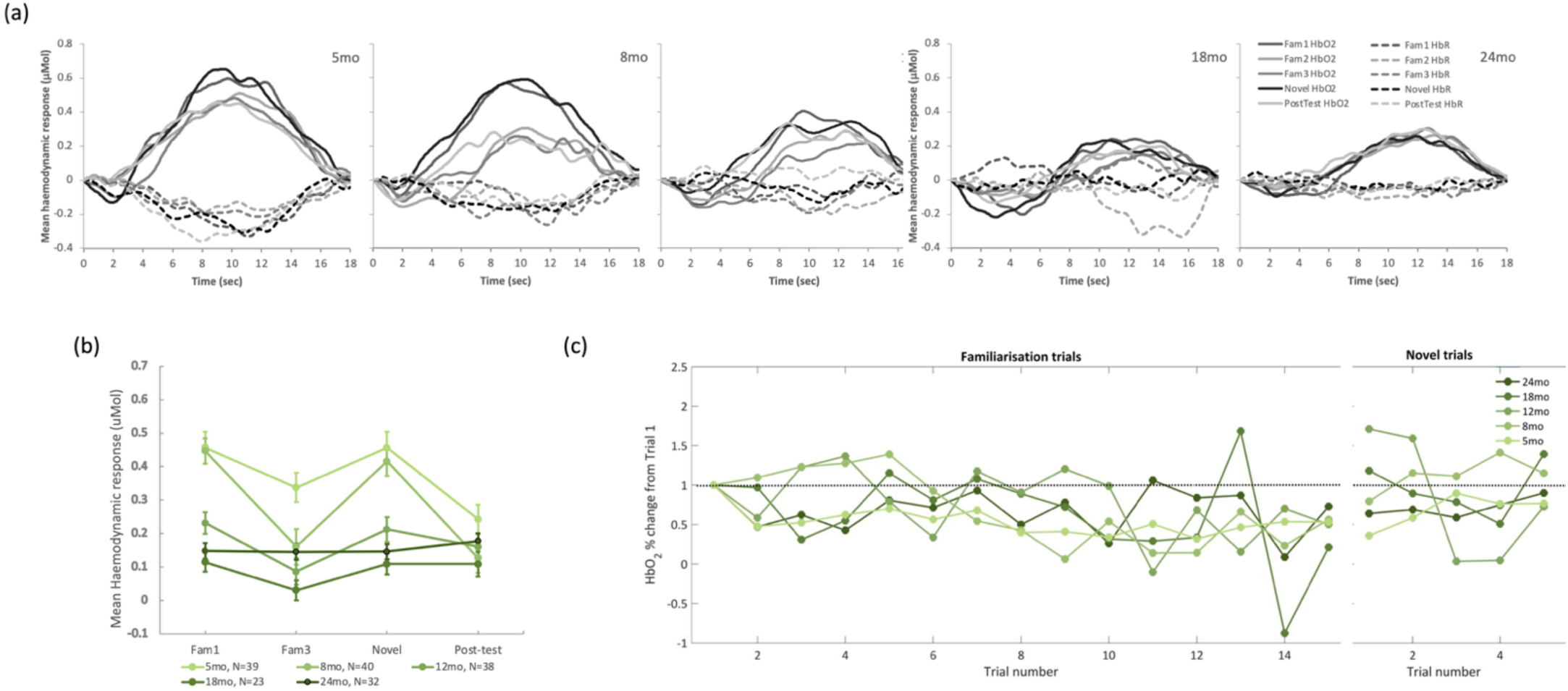
Cohort 2 (UK) Haemodynamic response. (a) Averaged HbO2 (solid lines) and HbR (dashed lines) across all participants at each age point; different epochs are represented by shades of grey. (b) Average HbO2 change from baseline within a time window post stimulus onset (8 to 12sec for 5 and 8mo; 10 to 14 for 12 to 24mo) across valid trials per epoch, in % change from Fam1 at each age point (represented in shades of green as specified in the legend in Fig3.c). (c) Trial by trial average HbO2 change from baseline within a time window post stimulus onset (8 to 12sec for 5 and 8mo; 10 to 14 for 12 to 24mo), for all familiarization trials (1 to 15) and Novel (16-20) epochs.

**Table 4.** Epoch effect within each visit, UK cohort. General Linear model with repeated measures (Epoch), including post-hoc pairwise comparisons between the most relevant epochs, Bonferroni-corrected for multiple comparisons.

| Visit | N | Epoch effect |  |  | Significant Pairwise comparisons (p-values, Bonferroni-corrected) |  |  |  |
| --- | --- | --- | --- | --- | --- | --- | --- | --- |
|  |  | F | p-val | h2 | Fam1>Fam3 | Fam1 > Novel | Fam1>Post-test | Novel>Fam3 |
| 5mo | 39 | 7.964 | 0.008 | 0.199 |  |  |  | 0.006 |
| 8mo | 40 | 15.229 | <0.001 | 0.286 | 0.005 |  | 0.001 | 0.032 |
| 12mo | 38 | 4.51 | 0.042 | 0.127 |  |  |  |  |
| 18mo | 23 | 1.561 | 0.225 | 0.066 |  |  |  |  |
| 24mo | 33 | 0.404 | 0.75 | 0.014 |  |  |  |  |

At 18mo, one outlier was identified at Fam1. By removing this outlier, Epoch effect remains not significant (F=1.190; p=0.177; partial eta square=0.085)

At 18mo, infants’ responses to the task became weaker and disappeared at 24mo (Figure 5a and 5b). At both visits sphericity was met (18mo: Χ^2^=3.687, p=0.596;24mo: Χ^2^=5.2, p=0.362). At 18mo, the number of datasets included in the analysis at 18mo was lower than at all other visits, with higher exclusion rates for fussing out and headgear placement (Figure 1).

Finally, linear mixed models with visit as a repeated effect to investigate the effect of age on the response to HaND revealed a significant age effect on Habituation (F[4,59.594] =3.344; p=0.015) and on Novelty Detection (F[4,59.118]=2.597; p=0.045).

### 3.2 Threshold-Free Cluster Enhancement (TFCE)

The results of the TFCE analysis are represented in Figures 6 (GM) and 8 (UK), for the epoch vs baseline statistics; and Figures 7 (GM) and 9 (UK) for the epoch contrasts. ROI locations with activation spatial maps corresponding to TFCE statistics for the Fam1 epoch are presented in Figures S1 and S2 of the Supplementary Information. Graphical representations of the haemodynamic responses for Fam1, Fam3, and Novel conditions—including periods showing significant contrasts across all time points and in both cohorts—are available in Figures S3 to S12 of the Supplementary Information.

**Figure 6.**
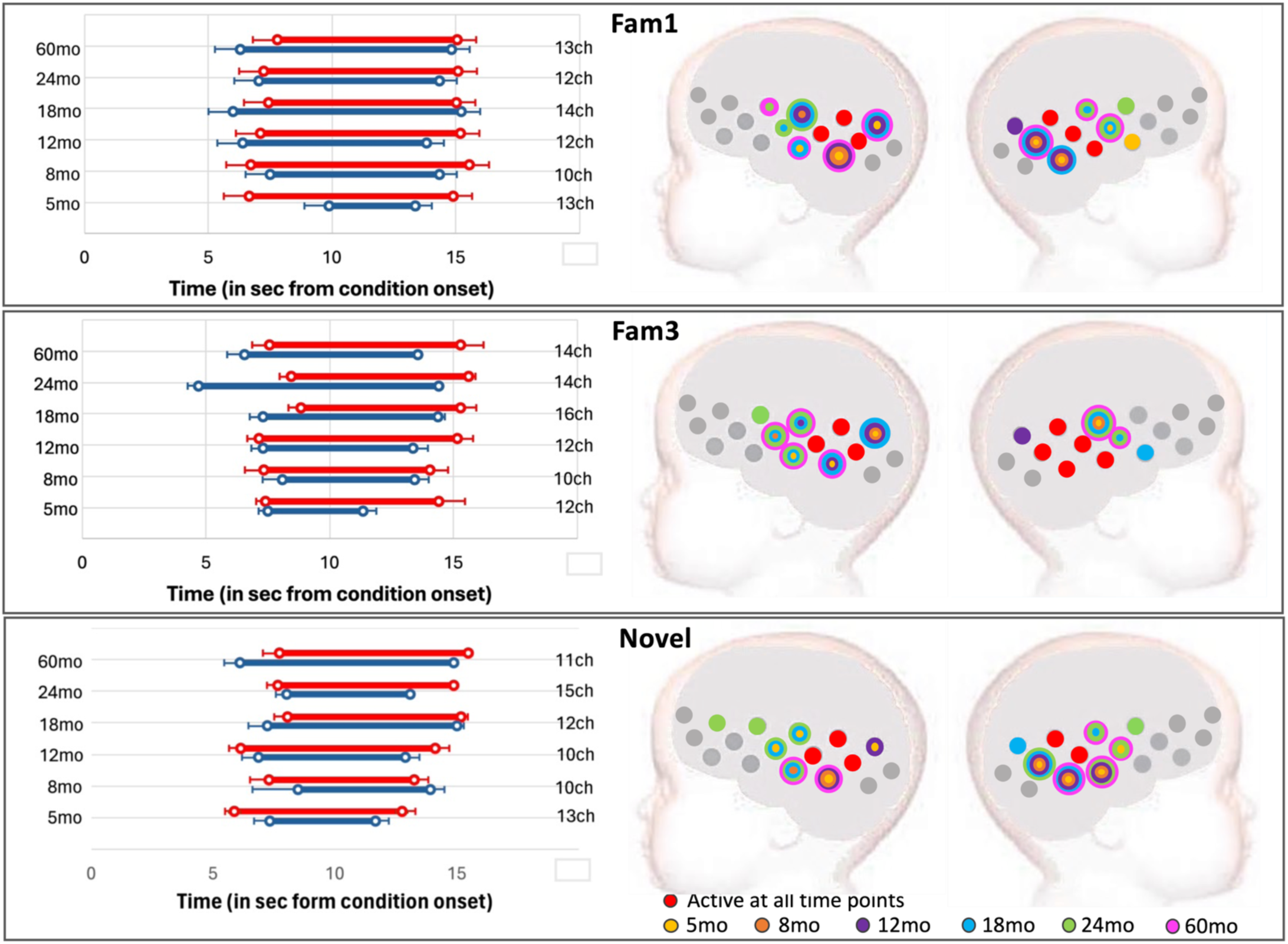
GM cohort: activation at epochs Fam1, Fam3 and Novel. Left: Averaged time window of significant increase in HbO2 (red) and/or HbR (blue) consistent with TFCE statistics; error bars indicate standard error of the mean; the vertical axis has visit (left) and number of active channels (right). Right: spatial mapping of activation consistent with TFCE statistics, color-coded by visit, considering HbO2 increase and HbR decrease.

**Figure 7.**
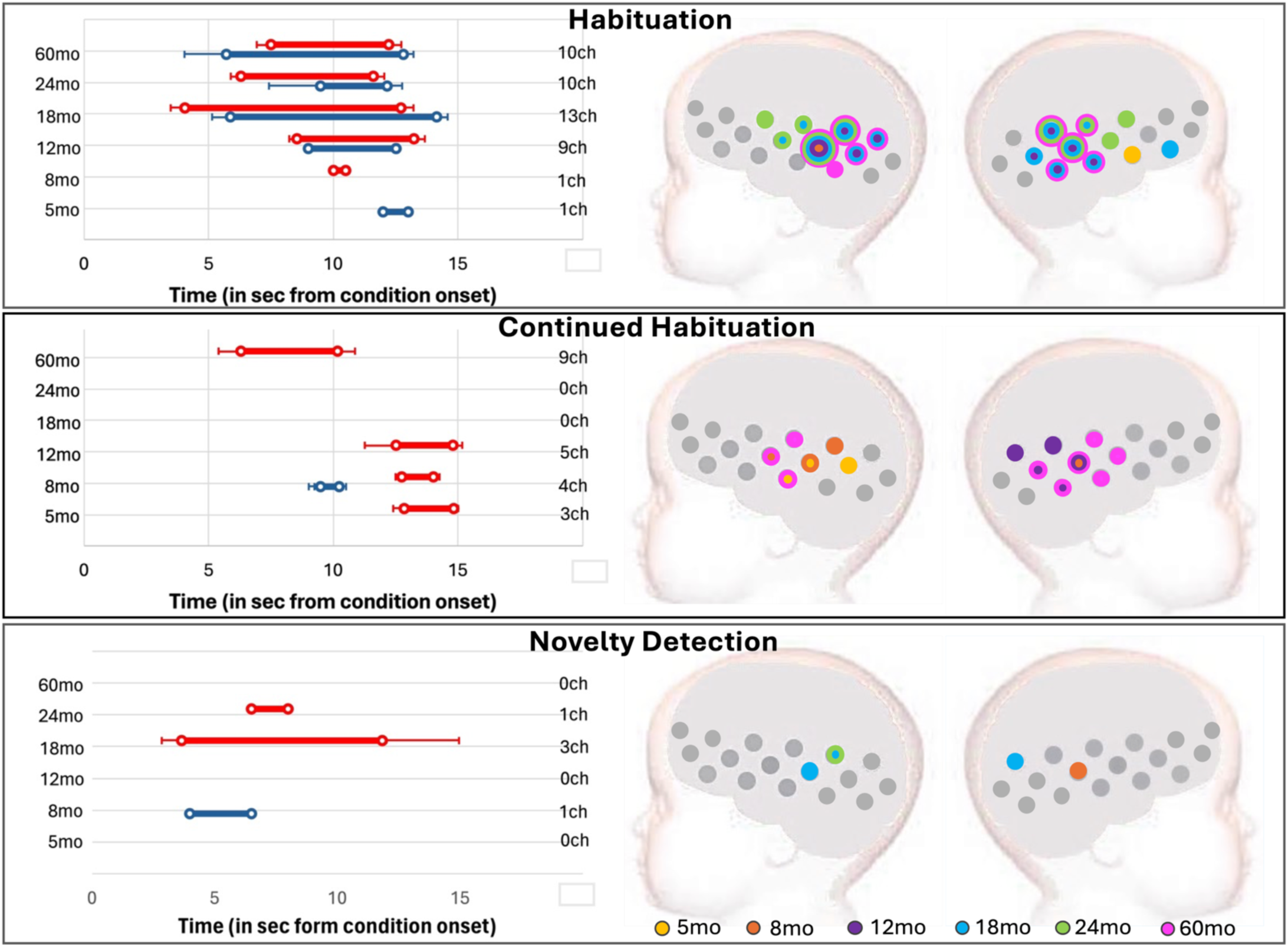
GM cohort: Left: Right: Averaged time window of significant epoch contrast in HbO2 (red) and/or HbR (blue) consistent with TFCE statistics. Right: Spatial mapping of significant epoch contrasts consistent with TFCE statistics considering HbO2 and HbR: Habituation (Fam1>Fam3); Novelty detection (Novel>Fam3); and Continued habituation (Fam3>Novel), color-coded by visit.

#### 3.2.1 Cohort 1 (GM)

HaND responses in the Gambian cohort were characterized by large clusters: at Fam1, clusters included from 10 active channels at 8mo, to 14ch at 18mo; at Fam3, from 10ch at 8mo to 16ch at 18mo; and at Novel, from 10ch at 8 and 12mo, to 15ch at 24mo. Significant Fam3 activation was present at all visits, and Continued Habituation (Fam3>Novel) was observed at 5, 8, 12 and 60mo. Interestingly, the TFCE analysis also revealed Novelty Detection in a few channels at 18 and 24mo (Figures 6 and 7).

The clusters that characterized activation at each epoch largely overlapped across visits (Figure 6), and 3 channels on the left hemisphere and 2 on the right led activation across all visits and epochs. Timing of the responses remained stable across chromophores, visits and epochs and were consistent with the expected haemodynamic response to the conditions (see Figure 6, left panels).

The spatial map of activation was consistent across the two analysis methods: (1) the ROIs calculated with the CPA overlapped with the clusters found with the TFCE at all visits (see Figure S1 of the Supplementary Information); and (2) the temporal windows used with the CPA method (8 to 12sec post stimulus onset at *5* and 8mo; 10 to 14 at the remaining visits), were within the temporal boundaries of the TFCE-calculated clusters.

Epoch contrasts revealed clusters driving Habituation at all visits (figure 7, upper panel), starting at 5 and 8mo with 1-channel clusters. Habituation clusters became larger at older ages, with 9 channels at 12mo, 13 at 18mo, 10 channels at 24 and 60mo. A continued decrease of the signal beyond the initial familiarization period (Continued Habituation, figure 7, middle panel) was detected at 5mo, in a 3-channel cluster (HbO2 only); 8mo, in a 4-channel cluster (HbO2 and HbR); 12mo, in a 5-channel cluster (HbO2 only); and 60mo, in a 9-channel cluster (HbO2 only). The TFCE analysis provided increased sensitivity for the Novelty Detection contrast, by considering all available channels and for not restricting the analysis to a pre-defined time window. In this case, Novelty Detection was driven by a 3-channel cluster at 18mo and a 1-channel cluster at 24mo. The single channel with Novelty Detection at 8mo should be interpreted with caution: the HbR signal in this channel showed Novel>Fam3 between 4 and 6.5sec post stimulus onset, while the HbO2 signal showed Fam3>Novel between 12.5 and 14sec. Given that the evidence is stronger for Fam3>Novel at this age (as listed above), it is reasonable to consider this as a false positive.

#### 3.2.2 Cohort 2 (UK)

In the UK, activation at Fam1 was led by clusters that decreased in size with age: 11 channels at 5mo; 11ch at 8mo; 5ch at 12mo, 1ch at 18mo and 24mo (see Figure 8). The ROIs calculated in section 3.1.2 overlapped these clusters (Figure S2 of the Supplementary Information). Temporally, Fam1 activation started at increasingly later time with age: at 5mo, activation was led by a cluster starting at 7.4sec and 10.2sec post-stimulus onset (for HbO2 and HbR, respectively); at 8mo, starting at 7.7 and 11.2sec; at 12mo, starting at 8.6sec and 14sec; at 18mo, starting at 13.5sec (HbO2 only); and at 24mo, starting 14sec (HbO2 only). Fam3 activation was led by smaller clusters, with onset times less consistent than for Fam1 (Figure 8 middle panel). A 6-channel cluster led Fam3 activation at 5mo, starting at 8.4 and 8.3sec post-stimulus onset (HbO2 and HbR, respectively); at 8mo, by a 2-channel cluster starting at 9.7sec (HbR only); at 12mo, by a 2-channel cluster starting at 7.7sec (HbO2 and HbR); at 24mo, by a 7-channel cluster starting at 13.3sec and 7.5sec post condition onset. No Fam3 activation was detected at 18mo. Novel activation was led by clusters that increased in size from Fam3 (Figure 8, lower panel): at 5mo, 11ch starting at 7.9 and 7.6sec; at 8mo, 11ch starting at 8sec (HbO2); at 12mo, 8ch starting at 11.1sec (HbO2); and at 24mo, by 3ch starting at 10.1sec.

**Figure 8.**
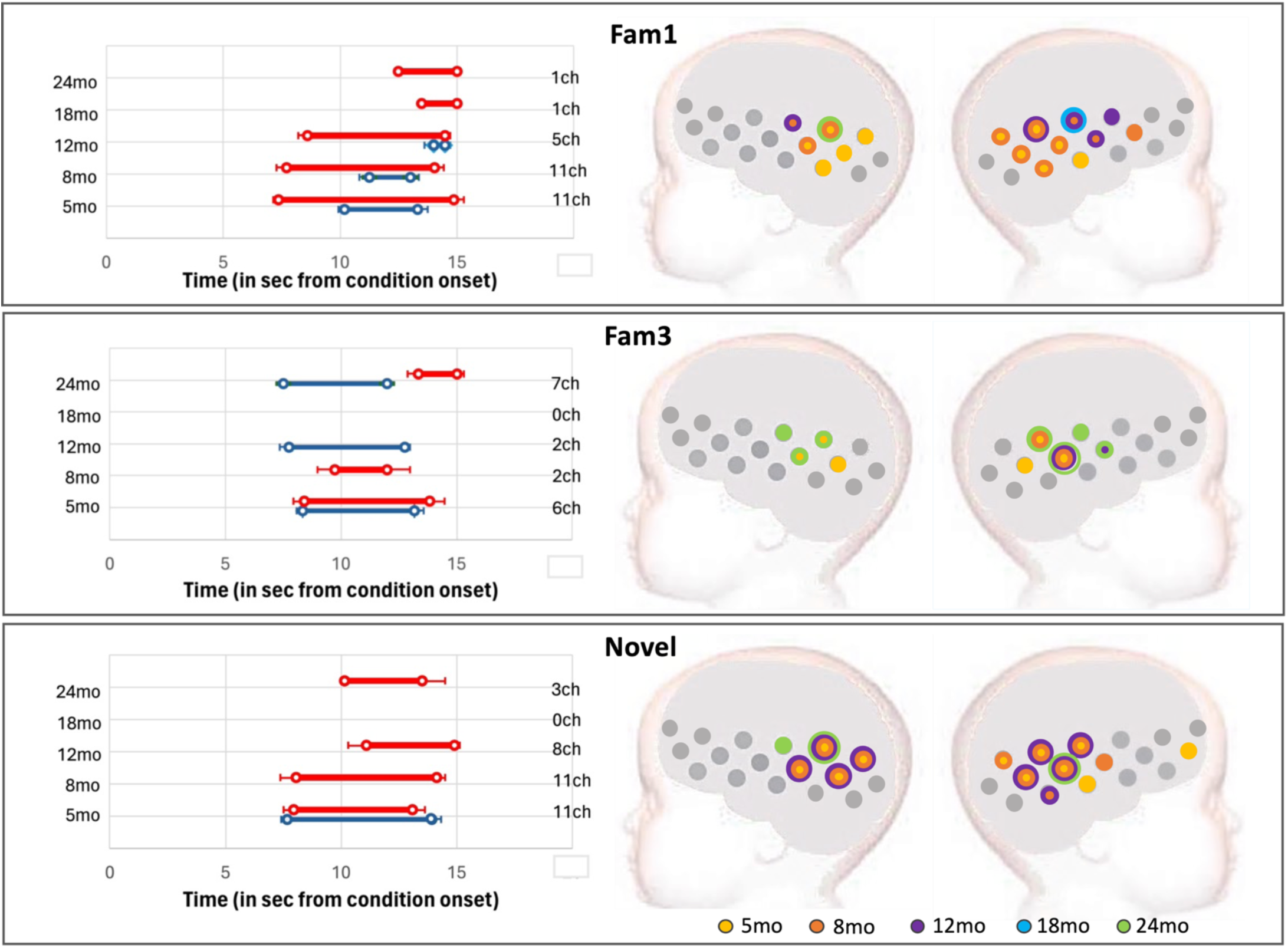
UK cohort: activation at epochs Fam1, Fam3 and Novel. Left: Averaged time window of significant increase in HbO2 (red) and/or HbR (blue) consistent with TFCE statistics; error bars indicate standard error of the mean; the vertical axis has visit (left) and number of active channels (right). Right: spatial mapping of activation consistent with TFCE statistics, color-coded by visit, considering HbO2 increase and HbR decrease.

TFCE analysis of the epoch contrasts revealed significant Habituation at 8 and 12mo (Figure 9, top panel). At 8mo, the clusters included 4 channels, and at 12mo, 1 channel (HbO2 only in both cases). The Novel vs Fam3 contrast did not show Continued Habituation in this cohort (Figure 9, middle panel), however, in line with the results from section 3.1.2, it did reveal Novelty detection at the earlier visits (5, 8 and 12mo, Figure 9, lower panel). Novelty Detection at 5mo was led by a cluster that included two adjacent channels on the left hemisphere, starting at 11.5sec (HbO2) and 6sec (HbR) post stimulus onset; at 8mo, by a 4ch cluster starting at 6.4sec (HbO2 only); and at 12mo, by a 1ch cluster on the left hemisphere, starting at 6sec post-condition onset (HbO2). No Habituation or Novelty Detection were found at 18 or 24mo.

**Figure 9.**
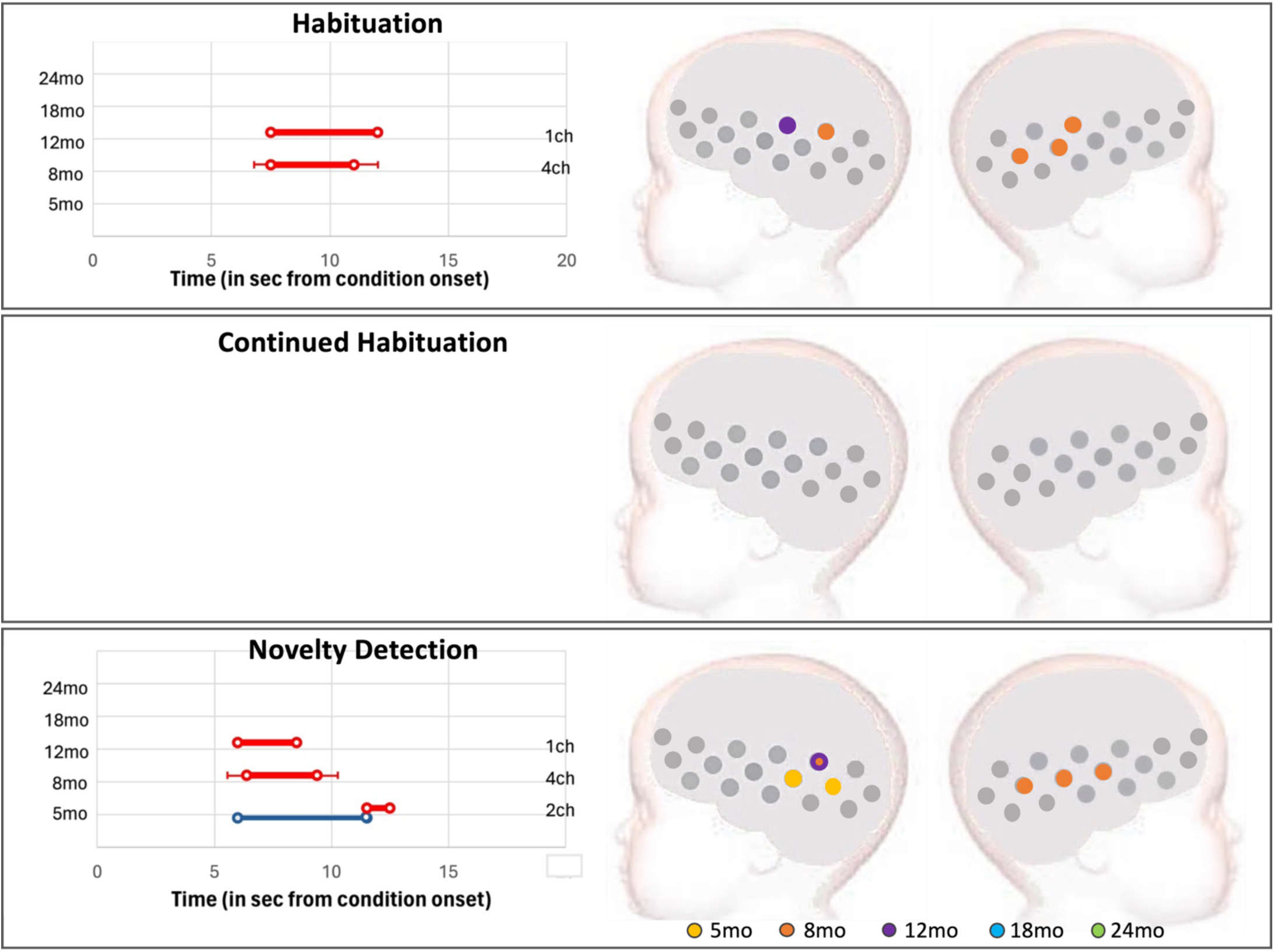
UK cohort: Left: Right: Averaged time window of significant epoch contrast in HbO2 (red) and/or HbR (blue) consistent with TFCE statistics. Right: Spatial mapping of significant epoch contrasts consistent with TFCE statistics considering HbO2 and HbR: Habituation (Fam1>Fam3); Novelty detection (Novel>Fam3); and Continued habituation (Fam3>Novel), color-coded by visit.

## 4. Discussion

This work describes (a) the haemodynamic response to a repeated presentation of identical auditory stimuli and (b) the response when a novel stimulus is introduced, in two cohorts of infants from The Gambia and the UK. We report longitudinal results across 5, 8, 12, 18 and 24 months of age, with an additional visit for the Gambian cohort at 3 to 5 years of age (60mo). Here we follow up from our previous work where we reported results from both cohorts at 5 and 8mo, where the GM cohort included the first 100 infants studied only. We report results from the complete groups, with a longitudinal perspective of the change in the responses with age. Overall, while repeated stimulation elicits a decrease in the amplitude of the response (Habituation) in both cohorts, the presentation of the novel stimuli does not elicit a recovery of response (Novelty Detection) at all visits, in the GM cohort.

### 4.1 Summary of findings

**Cohort 1 (GM):** CPA analysis in the Gambian cohort found a significant epoch effect at all visits. More specifically, these results showed (1) evidence of Habituation at all visits, (2) a general trend of a continuation of habituation across the session at most visits, (3) and an apparent reversal of this trend with the presentation of the Novel stimuli at 18 and 24mo. At 5 and 8mo, even though Fam1>Fam3 was not significant, the amplitude of the response continued to significantly decrease from Fam1 to Novel and PostTest. However, at 18 and 24mo, there was an apparent interruption of the decreasing trend of the HbO2 amplitude with the presentation of the novel stimuli (Figure 4b and 4c). However, direct paired comparison of Fam3 vs Novel did not reach statistical significance.

Investigation with the TFCE analysis provided more insight into the temporal and spatial features of the responses. The analysis confirmed the existence of habituation during familiarization trials at all visits, including at 5 and 8mo, which had been overlooked by CPA. The increased sensitivity displayed by the TFCE approach was driven by (1) its freedom from the temporal (no fixed time window) and spatial (no fixed number of channels per cluster) constrains of CPA; and (2) we included both HbO2 and HbR signals. For example, at 5mo, in the GM cohort, Fam1>Fam3 was significant in the HbR signal only. In contrast with the CPA-based analysis, this approach also found evidence of Novelty Detection at 18 and 24mo - localized in 3 channels at 18mo and 1channel at 24mo. The fact that significant results were found in such small number of channels explains why this effect was missed in the analysis using the 6-channel CPA-defined ROIs: the signal was likely averaged out by activity in the other channels within the ROI.

Interestingly, similar patterns in the responses to familiar and novel conditions were reported across the BRIGHT study using a different experimental paradigm and imaging modality (EEG). At *1*, *5* and 18mo, participants also performed an auditory oddball task with EEG. Direct comparisons between fNIRS and EEG responses revealed (1) positive correlation between fNIRS and EEG indices of Habituation at 5mo and Novelty Detection at 5 and 18mo; and (2) significantly larger Habituation at 18mo relative to 5mo for both modalities(Katus et al. 2023).

**Cohort 2 (UK):** Infants in the UK showed an increasingly stronger HaND pattern with age between 5 and 8mo in agreement with our previous findings (Lloyd-Fox et al. 2019), and became weaker at 12mo. The pattern was still visible at 18mo, but it had become not significant at this point and continued this pattern at 24mo. The results provided by the two approaches were very similar although the TFCE provided further detail on the spatial and temporal localization of the response. Also, at 12mo this method provided evidence of Novelty Detection in this cohort, which the CPA-based analysis had missed. Similarly to the GM, this discrepancy may have been due to being very localized, thus it was potentially averaged-out by the activity in the other channels of the ROI.

### 4.2 How do the current findings relate to the wider literature?

The capacity to habituate to repeated stimulation is present from very early in development and, together with the response recovery that is observed after a change in stimulation, have been used in a number of studies that investigate language development using EEG (Woodruff Carr et al. 2021; Dehaene-Lambertz and Dehaene 1994) and fNIRS (Benavides-Varela and Gervain 2017; Gervain et al. 2008; Fló et al. 2019; Benavides-Varela et al. 2011; Bouchon, Nazzi, and Gervain 2015; Benavides-Varela et al. 2017) as it “provides a foundation for the learning and cognition on which higher functions are constructed and it could efficiently predict cognitive developmental outcomes in term and preterm newborn infants.” (Sicard-Cras et al. 2022).

Exposure to repeated stimuli can result in either repetition suppression or repetition enhancement in developmental populations, largely depending on the type of stimuli (Issard and Gervain 2017): repetition suppression (Habituation) is associated with the presentation of identical physical stimuli (Emberson et al. 2017; Nakano et al. 2009); whereas repetition enhancement has been reported in studies with repetition presented at a more structural level (such as sequences of repeated phonemes rather than identical sentences in each trial) (Gervain et al. 2008; Bouchon, Nazzi, and Gervain 2015). Our results (in both cohorts) are, overall, in line with the former, as we found a decreasing response with successive presentations of the same stimulation. However, many of these studies report responses in the frontal brain regions, which we did not consistently find here (frontal (IFG) activation to the Fam1 and Novel epochs was found in the GM cohort at 24months only). The overall lack of frontal response observed in both cohorts may be partially due to a discrepancy in the state of alertness/sleep with the study by Nakano et al. (Nakano et al. 2009), as Taga et al. found that, although auditory processing continues during sleep, brain responses are more localized during wakefulness (Taga, Watanabe, and Homae 2018). Frontal activation, however, was detected in the response to the Social task (in preparation), which was run before the HaND task in the fNIRS session. The strategy to engage the infants’ attention was different at each task: the Social task had auditory and visual stimuli, and no additional attention grabbers were needed; however, the HaND task required the visual distraction of the bubbles to engage the infant’s attention. The experimenters started the bubbles before starting the task and used them throughout. It is possible that the bubbles elicited a sustained response that masked frontal activation to the HaND sounds. Unfortunately, in the current data set we can’t test activation to the bubbles compared to a baseline segment as the time of onset of the bubbles was not recorded, and so we are unable to explore this potential impact. A further consideration is that the frontal regions may have shown greater variability in response between participants, which we aim to explore in future work where we explore responses at participant level in relation to contextual factors.

In contrast to our observations at younger ages, the response of the HaND profile weakened from 12mo onwards in the UK, and at 60mo in the GM, with a smaller amplitude in the signature of the response across epochs. A detailed analysis of risk and resilience factors in relation to the HaND response data and other domains of neurodevelopment assessed in BRIGHT will be the subject of future analyses. In the following paragraphs we discuss factors that may have impacted in the responses.

It is possible that that an increased presence of thick, long or curly hair with age may have interfered with the detection of the fNIRS signals. In the UK, 5.3% of the participants at 5mo had potentially problematic hair (long, thick and/or curly); this increased to 22.7% and 18.2% at *18* and 24mo respectively. In the same cohort, the amplitude of the response at Fam1 in decreased from 5mo to 24mo (with HbO2, paired t-test, t=3.040, p=0.006, Cohen’s d=0.85, see Figures 6a and 6b). However, this would not explain why the Gambian children at 60mo showed the strongest response to Fam1, yet their response to HaND returned to the Habituation-only pattern observed at younger ages (Figures 5a and 5b). Our results, however, resemble the findings from the longitudinal study by Deguire et al. (2022), where participants showed a stronger Novelty Detection at the first visit (6mo) than later visits (24mo and 48mo). Remarkably, these results were from Canadian children living in the Montreal area and align with our results from the UK cohort. While some Novelty Detection was present at 24mo Gambian infants also showed no Novelty Detection at 60mo. It is possible that, as Deguire et al (2022) argue, at infants may have outgrown the task, and it was no longer interesting enough to engage their attention in it.

Several factors may be behind the differences in the HaND signature between the two cohorts. It is possible that (1) the effects of Continued Habituation and minimal Novelty Detection in the GM, compared with the significant response to HaND until up to 12mo in the UK cohort may have been driven by the context in which these infants live. Although the aim of this work was to describe the developmental trajectories of the HaND response in the GM and UK infants and leave for future work the investigation of what factors may have influenced the responses, we will nevertheless mention potential external influences in these responses. Firstly, UK infants were from the area in and around Cambridge, a high-resource urban community, whereas GM participants resided in and around the village of Keneba, a low-resource rural community. Factors such as family and household arrangements may have shaped the infants’ exposure to sounds and social interaction which in turn could have influenced their responses to the HaND stimuli. From an early age, in this rural Gambian context, infants were typically carried on their mother’s back while the mothers engaged in daily tasks – such as farming and selling their produce at the local market. Additionally, the immediate home environment often included multiple generations living in large compounds (Hennig et al. 2017), meaning infants were routinely exposed to a high number and diversity of voices from birth. In contrast, the UK infants typically lived in more socially shielded environments, in smaller households, often looked after by a parent on leave. These differences in the daily environment may have shaped the infants’ ability to block out voices or sounds that were deemed less relevant.

Factors related to the low-resource Gambian region where the GM data was collected may also have impacted the responses. While not all our GM participants or their mothers and families may have faced adversity since conception, they would have experienced considerably higher rates of poverty-associated factors relative to the UK cohort. In the UK, none of the infants showed severe growth deficiency (defined as anthropometric z-scores lower than -3SD from WHO Median standards), and in The Gambia, very few infants did (3 infants at 18mo and 3 at 24mo). Using anthropometric z-scores < -2SD as an indicator of undernutrition, 18.8% of children at 18months of age were undernourished in The Gambia compared to only 8.7% in the UK. At 24months of age this comparison was 15.6% vs. 3.03%. In the critical period from birth to 24months, complex interactions exist between nutritional requirements and brain development and function. Maternal nutritional status, environmental exposures and inflammation can have a critical impact on neurodevelopment (Krebs, Lozoff, and Georgieff 2017). For instance, a significant correlation exists between early nutrient intake and brain maturation, and it is stronger in infancy and early childhood (between 6 and 60mo), (Schneider et al. 2023), which highlights the importance of implementing early interventions. Iron, an is an essential micronutrient related to cognitive and motor development, yet its deficiency is most prevalent in the first few years of life (Krebs, Lozoff, and Georgieff 2017). Other contextual factors such as food insecurity, infectious disease and psychological stress related to a child’s environment (Jensen, Berens, and Nelson 2017), as well as psychosocial and environmental factors associated with poverty (Drago et al. 2020; Hamadani et al. 2014) may shape a child’s developmental trajectory. For example, higher socio-economic status and the mother’s level of education have been linked to higher cognitive abilities in toddlers (Johnstone et al. 2021). This association was also observed in samples from rural India and Mid-Western USA, where poor maternal education and lower socio-economic status were correlated with weaker brain activity and poorer distractor suppression in a working memory task (Wijeakumar et al. 2019). Likewise, 6 and 36month-old infants living in an urban slum in Bangladesh showed a significant correlation between the magnitude of social selective responses and maternal stress, maternal education and the caregiving environment (Perdue et al. 2019), and further associations of social discrimination with exposure to psychosocial adversity (such as family conflict and maternal depression) were in infants recruited from the same population (Pirazzoli et al. 2022).

As part of the BRIGHT project, this data will contribute to further investigate modulation of contextual variables on cognitive development. The first outputs of this project have already been published. For example, the EEG protocol revealed associations with neurodevelopmental scores from the Mullen Scales of Early Learning (MSEL) at 5months of age across the two cohorts, suggesting that the neuroimaging markers possess a certain degree of universality that might make them suitable to serve as measures of neurocognitive development in a range of populations (Katus et al. 2020). In future work we aim to investigate the neural underpinnings of these differences by estimating the potential modulatory effects of context, and, for this, we will study correlations with behavioral measures such as parent-child interaction, and the Neonatal Assessment Behavioral Scale, as well as information about family context, environment and caregiving. Also, it will be interesting to explore whether responses to the HaND paradigm are predictors of outcomes later such as memory development, and thus if they are consistent with the observations from the EEG paradigm (Katus et al. 2022).

### 1.3 Strengths and Limitations

Co-registration with age-appropriate brain templates: Pictures of the infants wearing the fNIRS arrays before and after the session as well as anthropometric measurements from the UK cohort were used to investigate variability in head size and array position on the anatomical and statistical inferences drawn from the data(Collins-Jones et al. 2021). The study revealed that channel-space longitudinal analysis of fNIRS data, given the sample size, was robust to assumptions of head size and array position. In other words, the quality control protocol for reviewing fNIRS array placement on the head contributed to maintaining consistent headgear position across visits. The work identified the anatomical region of sensitivity for each channel of the array at each visit, from 1mo to 24mo (Table S1 of the Supplementary Information; the report included visits at *5*, *8* and 12mo) (Collins-Jones et al. 2021). Taking together these results and the strict protocol for headgear placement, based on alignment between anatomical landmarks ad reference points on the arrays (similar to (Blasi et al. 2014) and (Sarah Lloyd-Fox, Richards, et al. 2014), we are confident in the brain regions identified in this work for the UK cohort. An additional source of variability in the placement of the fNIRS headgear on the scalp is the size of the headband holding the arrays. The headband was designed to accommodate head growth, and the size used with each participant was selected at the beginning of each session based on measures of head circumference. Three headband sizes were available in the UK (small, medium and large) and two in The Gambia (small and large). Although great care was placed in consistently positioning the headgear on participants heads across visits and cohorts, it is possible that the decision on headband size to match the participants head circumference might have slightly influenced the sensitivity map of each cohort. Figure S14 of the Supplementary Information illustrates the use of each headband size per visit and cohort. Most infants in the GM cohort used the shortest headband at *5* and 8mo; and at 24mo, just over 25% of participants were still using this shortest headband. In the UK cohort however, at 5mo, 21% of participants were already using the longest headband, and 37% the medium one. At 12mo, 14% of participants were using the medium headband and the rest, the longest one; and from 18mo onward, the longest headband was used with all participants. Differences in the pace of head growth between the two cohorts drove the changes in use of each headband size, but the decision of which headband to use originated in the team of experimenters conducting the sessions, therefore there is a chance of small variability between sites of array placement. Here we have assumed that the regions identified in (Collins-Jones et al. 2021) were identical in both cohorts.

Comparisons between the two cohorts: This study was designed to describe the development of the brain responses to repeated stimuli, and to a change in the form of a change in speaker in two cohorts, as infants in each cohort developed in different contexts. The study was not designed to present direct comparisons between the two. The UK cohort provided a more comprehensive longitudinal dataset to align with broader results from research studies in more extensively researched high-income populations. However, due to lack of previous research in similar African or LMIC contexts, by necessity the UK results and previously published data from studies using similar paradigms informed the hypothesis for the Gambian cohort and influenced our reflections on the findings.

Choice of stimulus presentation: At the time of the data collection, electricity was not available in the households of the West Kiang region of The Gambia, where most families in this cohort lived, therefore, contrary to infants in the UK, it is unlikely that GM infants had had prior exposure to non-live speech (i.e. Television, screen-based videos). Previous research has demonstrated that infants’ brain responses are significantly different when observing action in a person vs an object in motion in a live setting, but not when the stimuli are presented in pre-recorded videos (Shimada and Hiraki 2006). Furthermore, the ability to discriminate sounds within language is easier for infants when presented live relative to audio only (Kuhl, Tsao, and Liu 2003). These results support the push for experimental paradigms with ecologically valid designs with bidirectional interaction and contingent responsiveness that minimize the effects that “artificial” designs might induce in the brain responses (Gervain et al. 2023). However, in the current study this was not possible, and the extent of this effect on our results is unknown.

Multilingualism: In both cohorts some families were multilingual and the magnitude of this effect on the infant’s ability to discriminate between the different speakers is yet unexplored. In the GM while all families were Mandinka, for some infants some of their caregivers spoke a second language and may have used this in their presence. This has not been fully characterized for this cohort as the information was not available at the time of data collection. In the UK approximately one third of the families were multilingual, with some infants being exposed to a low level of English (see (Sarah Lloyd-Fox et al. 2024)), or differing levels of English depending on age point and who was providing the caregiving (i.e. parent/family member/childcare).

Characteristics of the stimuli between cohorts: while the female and male voices speaking the sentence in the experimental sessions in the UK and the female voice (familiarization) in the GM had similar characteristics of *pitch* and *intensity*, the range of these in the male GM voice was narrower. A crude analysis of the sounds indicated that the minimum *pitch* (=77.52) of the male GM voice fell within the mean ±2 standard deviations of the other three sounds (=97.54±21.80); but the mean (=172.88) and maximum (=252.76) *pitch* were below the other sounds (mean pitch=230.66±38.19; max pitch=477.60±57.22). Minimum *intensity* of the male GM voice (=48.70) was higher than the mean ±2 standard deviations of the other three sounds (=30.56±7.75) but mean (=73.95) and maximum (=83.63) *intensity* of the male GM voice were within mean ± 2 standard deviations of the other three sounds (=75.18±2.40; and 85.55±4.40). Therefore, we can’t completely rule out the possibility that the less salient male sound in the GM may have elicited different responses to the Novel stimuli relative to the stimuli contrast in the UK. Actors recording their voices for the experimental sessions were not given instructions to follow a specific intonation pattern, but rather to speak in a natural way as they would do to babies. While that would have made the stimuli more similar across sites, it might have sounded strange for the Gambian infants, as it would have altered the way a typical male voice sounds in their familiar context.

Exposure to adversity: Brain responses in the GM infants may have been conditioned by exposure to adversity since conception, but we don’t know the extent of this influence yet. Further analysis using correlations and models of habituation and Novelty Detection and markers of nutrition, psychosocial interaction, behavior, etc., will allow us to find the factors with the most influence on brain development.

Physical characteristics: The presence of hair in general can have a negative effect on the quality of data by partially or completely obstructing the path of light travelling from source to detectors via skin scalp and the cortex, and by making the headgear unstable and creating motion artefacts. A qualitative assessment of the presence of hair on each participant was recorded at each session. In the UK, 8% of participants included in the analysis had “long”, “a lot” or “thick” hair at 5 and 8mo, this percentage rose to 22% and 16% at 18 and 24mo, respectively; only one participant at 5mo and a different one at 8mo had “no hair”. It is customary in The Gambia to shave the infant’s head at 7 days of age, and to continue to shave the boys’ heads during early life. While 17% of infants at 5mo and 20% at 8mo had “thick”, “long”, “afro” or “rough” hair, at 18 and 24mo these numbers were as low as 4% and 10%. Moreover, in the GM, over 30% of infants at all visits had a shaved head, and the percentages rose to 65% at 12mo, 73% at 18mo and 67% at 24mo. The high percentage of infants with a shaved head is likely to be one of the leading reasons for the good quality of the data in the GM cohort. Figure S15 in the Supplementary Information illustrates the haemodynamic response at 5, 8, 18 and 24mo in the UK cohort in channel 12, a channel recording from STG at all visits that was included in all ROIs calculated with CPA. At 5 and 8mo, the haemodynamic response was significant and clearly visible, whereas at 18 and 24mo it was not.

## Conclusion

This work presents the first longitudinal study of the response suppression (habituation) and response recovery (Novelty Detection) from 5months of age to early toddlerhood in two cohorts of infants, one living in a high-resource setting typical of the majority of research being published to date; and a second cohort living in a low-resource setting in sub-Saharan Africa, representing a population that until recently was beyond the reach of functional brain imaging research. The Gambian infants’ response was characterized predominantly by Habituation, with weak Novelty Detection at 18 and 24mo; in the UK, a clear pattern of Habituation followed by Novelty detection was observed. These different patterns highlight the need to investigate at participant level the effects of socio-economic, health, environmental as well as cognitive and developmental markers on the responses. This information will contribute to our understanding of the factors that modulate developmental trajectories, which, in turn, may help formulate intervention strategies and evaluate their efficacy.

## Data availability

### Underlying data

The data used to support this study are stored in the Brain Imaging for Global Health Data Repository. The conditions of our ethics approval do not allow public archiving of pseudo-anonymized study data. The data cannot be fully anonymized due to the nature of combined sources of information, such as neuroimaging, sociodemographic, geographic and health measures, making it possible to attribute data to specific individuals, and hence, falling under personal information, the release of which would not be compliant with GDPR guidelines unless additional participant consent forms are completed. Access to any data collected during or generated by the BRIGHT project is fully audited, and to ensure data security, is overseen by the data management team in the UK and The Gambia. Our data sharing procedures were created in consultation with stakeholders and external consultation (Begum-Ali et al., 2023). To access the data, interested readers should contact the BRIGHT coordinator on the Contact page of our website. Access will be granted to named individuals following ethical procedures governing the reuse of sensitive data. Specifically, requestors must pre-register their proposal and clearly explain the purpose of the analysis to ensure that the purpose and nature of the research is consistent with that to which participating families originally consented. Additionally, requestors must complete and sign a data sharing agreement to ensure data is stored securely. Approved projects would need to adhere to the BRIGHT project’s policies on Ethics, Data Sharing, Authorship and Publication.

## Grant Information

The BRIGHT Study was funded by the Bill and Melinda Gates Foundation (OPP1127625) and core funding MC-A760-5QX00 to the International Nutrition Group by the Medical Research Council UK and the UK Department for International Development (DfID) under the MRC/DfID Concordat agreement. Lloyd-Fox and Blanco were further supported by a Medical Research Council Programme Grant MC-A760-5QX00 and a UKRI Future Leaders fellowship (grant MR/S018425/1). Milosavljevic was supported by an ESRC secondary Data Analysis Initiative Grant (ES/V016601/1).

## Acknowledgements

We are indebted to the parents and infants who took part in this study without whom this work would not have been possible. We also thank the wider team of staff at the MRC Keneba Field Station for supporting us in the collection of this data.

The authors would like to thank Professor Judit Gervain (University of Padova) for the analysis of the characteristics of the voice stimuli used in the experiments, and for her advice and interpretation of these.

## Notes

### Competing Interest Statement

The authors have declared no competing interest.

### Summary of Updates

Add a co-author that had been omitted in the previous version bt mistake

