## Supplementary information for "Longitudinal Habituation and Novelty Detection neural responses from infancy to early childhood in The Gambia and UK"

### Supplemental Information

| Hemisphere | Lobe | Region | 5mo | 8mo | 12mo | 18mo | 24mo |
|---|---|---|---|---|---|---|---|
| LEFT | FRONTAL | IFG | 3,5,7,8 | 3,5,7,8 | 3,5,7,8 | 3,5,7,8 | 3,5,7,8 |
| | | MFG | 2 | 2 | 2 | 2 | 2 |
| | | pre/postcentral gyrus | 6 | 6 | 6,14 | 6 | 6,14 |
| | TEMPORAL | ITG | 15 | 15 | 15 | 15 | |
| | | MTG | 9,13,16,17,18,19 | 9,13,16,17,18,19 | 9,13,16,17,18,19 | 9,13,16,17,18,19 | 9,13,15,16,17,18,19 |
| | | STG | 10,11,12,14 | 10,11,12,14 | 10,11,12 | 10,11,12,14 | 10,11,12 |
| RIGHT | FRONTAL | IFG | 22,26,27 | 22,26,27 | 22,25,26 | 22,25,26,27 | 22,25,26,27 |
| | | MFG | 20,23 | 20,23 | 20,23,27 | 20,23 | 20,23 |
| | | pre/postcentral gyrus | 25 | 25,33 | 33 | | |
| | TEMPORAL | ITG | 34,37 | 34,37 | 34 | 34 | 34 |
| | | MTG | 28,35,36,38 | 28,35,36,38 | 28,35,36,37,38 | 35,36,37,38 | 28,35,36,37,38 |
| | | STG | 29,30,31,32,33 | 29,30,31,32 | 28,29,30,31,32 | 28,29,30,31,32,33 | 28,29,30,31,32,33 |

Table S1. Brain regions identified with the co-registration of the fNIRs channels, (Collins-Jones et al. 2021).

**Fam1**

| | Channels LH | | | t-val | p-val | Channels RH | | | t-val | p-val |
|---|---|---|---|---|---|---|---|---|---|---|
| **5mo** | 12 | 16 | 13 | 22.1697 | <0.001 | 32 | 35 | 31 | 21.5874 | 0.001 |
| **8mo** | 12 | 16 | 13 | 21.2634 | <0.001 | 32 | 35 | 31 | 22.6406 | <0.001 |
| **12mo** | 12 | 16 | 13 | 28.3883 | 0.001 | 32 | 35 | 31 | 33.0329 | <0.001 |
| **18mo** | 12 | 13 | 14 | 27.7554 | 0.0018 | 32 | 35 | 31 | 29.0743 | 0.0026 |
| **24mo** | 12 | 13 | 14 | 23.2788 | 0.002 | 31 | 32 | 33 | 25.2869 | 0.002 |
| **60mo** | 12 | 16 | 13 | 48.9452 | 0.003 | 28 | 32 | 33 | 45.9056 | 0.004 |

**Novel**

| | Channels LH | | | t-val | p-val | Channels RH | | | t-val | p-val |
|---|---|---|---|---|---|---|---|---|---|---|
| **5mo** | 12 | 16 | 13 | 22.1697 | 0.0012 | 32 | 35 | 31 | 21.5875 | 0.0018 |
| **8mo** | 12 | 16 | 13 | 21.2634 | 0.004 | 32 | 35 | 31 | 22.6406 | 0.001 |
| **12mo** | 12 | 16 | 13 | 17.3536 | 0.001 | 32 | 35 | 31 | 16.0762 | 0.002 |
| **18mo** | 12 | 13 | 14 | 20.7611 | 0.004 | 32 | 35 | 31 | 19.0226 | 0.004 |
| **24mo** | 12 | 13 | 14 | 18.8013 | 0.005 | 31 | 32 | 33 | 23.4791 | <0.001 |
| **60mo** | 12 | 16 | 13 | 36.5880 | 0.0082 | 28 | 32 | 33 | 40.0536 | 0.0048 |

Table S2. GM: Cluster Permutation results for ROI selection with Fam1 and Novel epochs. RH = Right Hemisphere; LH = Left Hemisphere.

| | Fam1 | | | | | | | | | |
|---|---|---|---|---|---|---|---|---|---|---|
| | Channels LH | | | t-val | p-val | Channels RH | | | t-val | p-val |
| **5mo** | 12 | 16 | 13 | 12.3122 | 0.0008 | 32 | 35 | 31 | 11.5454 | 0.0042 |
| **8mo** | 12 | 13 | 14 | 11.8912 | 0.006 | 32 | 35 | 31 | 15.1159 | 0.001 |
| **12mo** | 12 | 13 | 14 | 7.9262 | 0.0402 | 29 | 32 | 33 | 0.0024 | 0.0058 |
| **18mo** | 12 | 13 | 14 | 3.4588 | 0.0362 | 29 | 32 | 33 | 3.3364 | 0.0014 |
| **34mo** | 12 | 13 | 14 | 7.9045 | 0.001 | 29 | 32 | 33 | 7.3036 | 0.003 |

| | Novel | | | | | | | | | |
|---|---|---|---|---|---|---|---|---|---|---|
| | Channels LH | | | t-val | p-val | Channels RH | | | t-val | p-val |
| **5mo** | 12 | 16 | 13 | 13.2818 | 0.003 | 32 | 35 | 31 | 11.4466 | 0.005 |
| **8mo** | 19 | 16 | 12 | 9.0786 | 0.017 | 32 | 35 | 31 | 12.8245 | <0.001 |
| **12mo** | 12 | 16 | 13 | 7.8008 | 0.002 | 31 | 32 | 33 | 8.1528 | 0.001 |
| **18mo** | 12 | 13 | 14 | 3.4156 | 0.009 | 29 | 32 | 33 | 3.4258 | 0.011 |
| **24mo** | 12 | 13 | 14 | 7.7284 | <0.001 | 29 | 32 | 33 | 6.6413 | 0.015 |

Table S3. UK: Cluster Permutation results for ROI selection with Fam1 and Novel epochs. RH = Right Hemisphere; LH = Left Hemisphere.

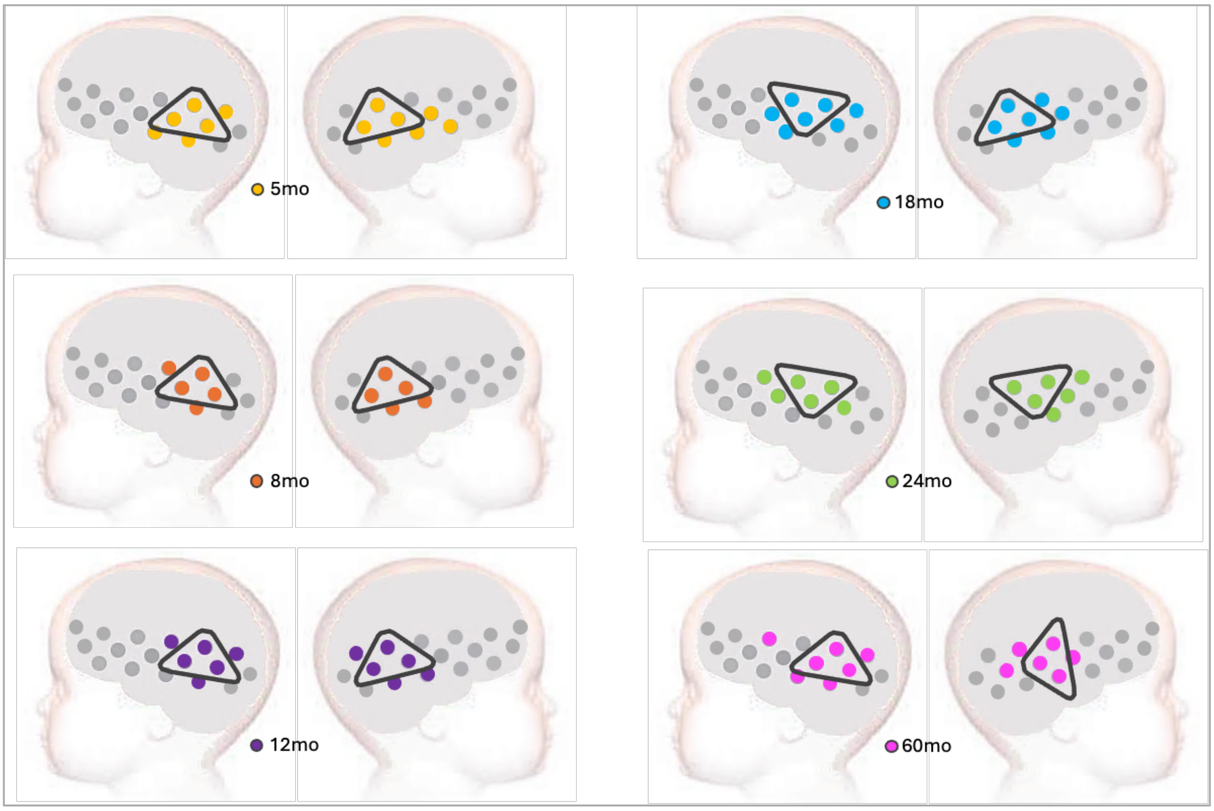


Figure S1. GM: Spatial mapping of significant activation at Fam1 consistent with TFCE statistics considering HbO2 and HbR. The triangular shapes represent the ROIs from the CPA.

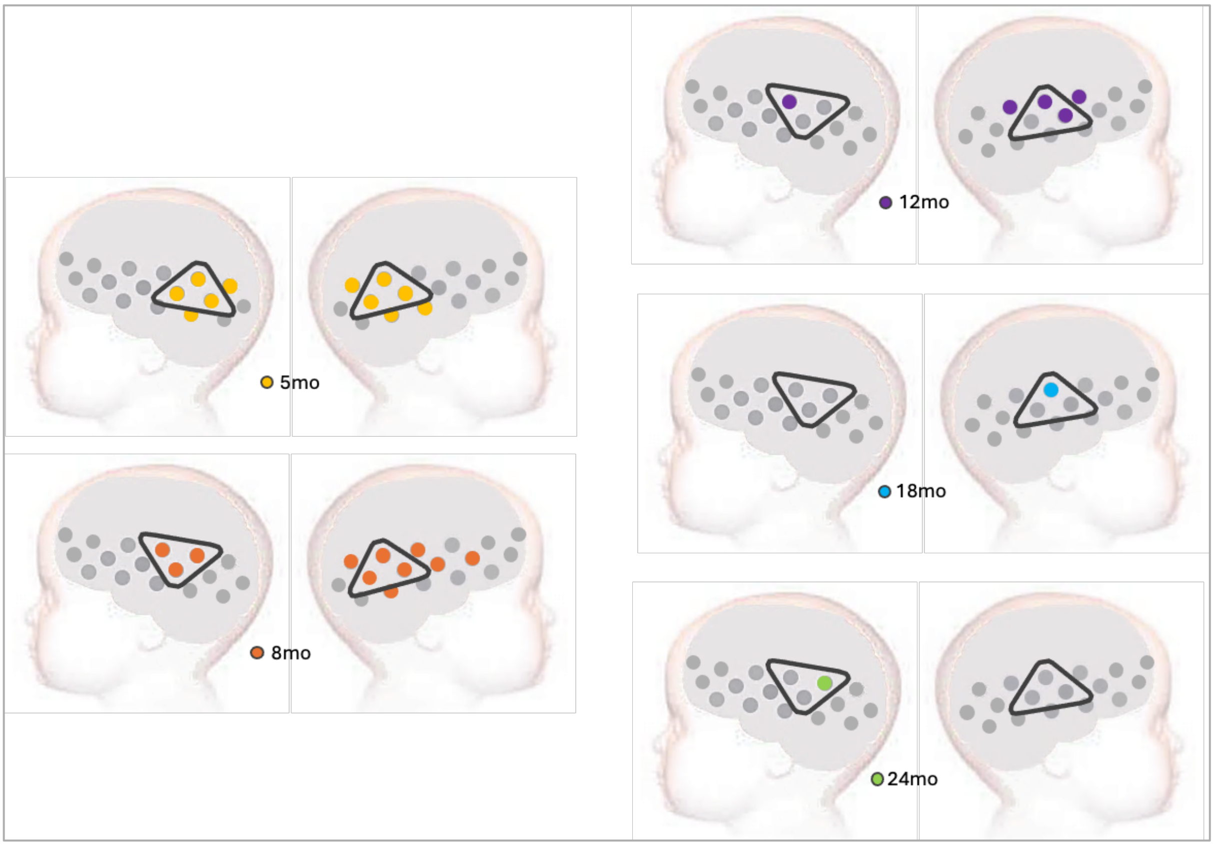


Figure S2. UK: Spatial mapping of significant activation at Fam1 consistent with TFCE statistics considering HbO2 and HbR. The triangular shapes represent the ROIs from the CPA.

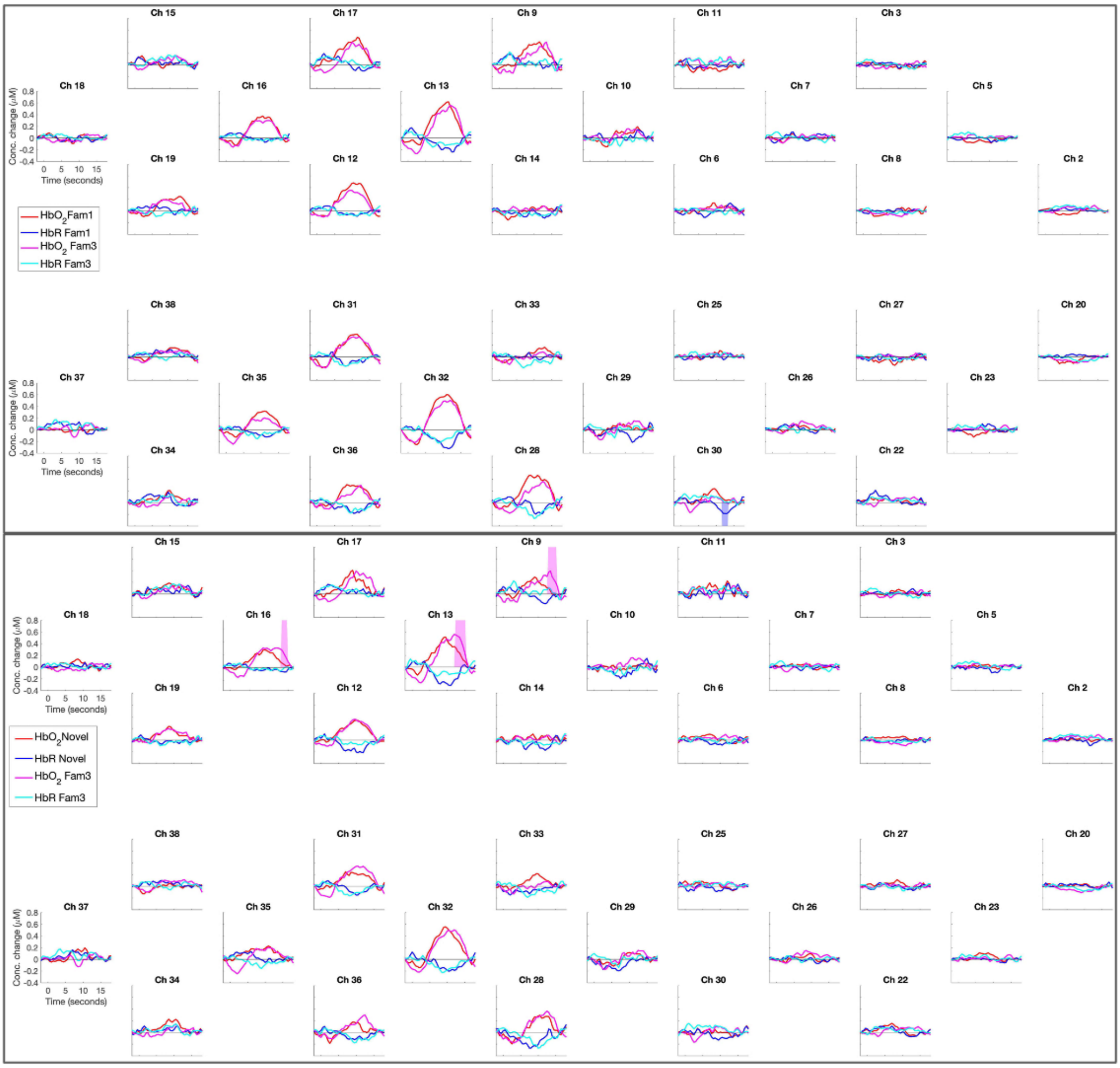


Figure S3. GM, 5mo visit: time courses of Fam1, Fam3 and Novel epochs. Upper panel: Fam1 vs Fam3 contrast. Lower panel: Novel vs Fm3 contrast. Shaded areas indicate significant difference between contrasts, the colour indicates which epoch is stronger (i.e. in the lower panel, magenta represents Fam3 HbO, then the magenta shaded areas represent Fam3 response is stronger than Novel).

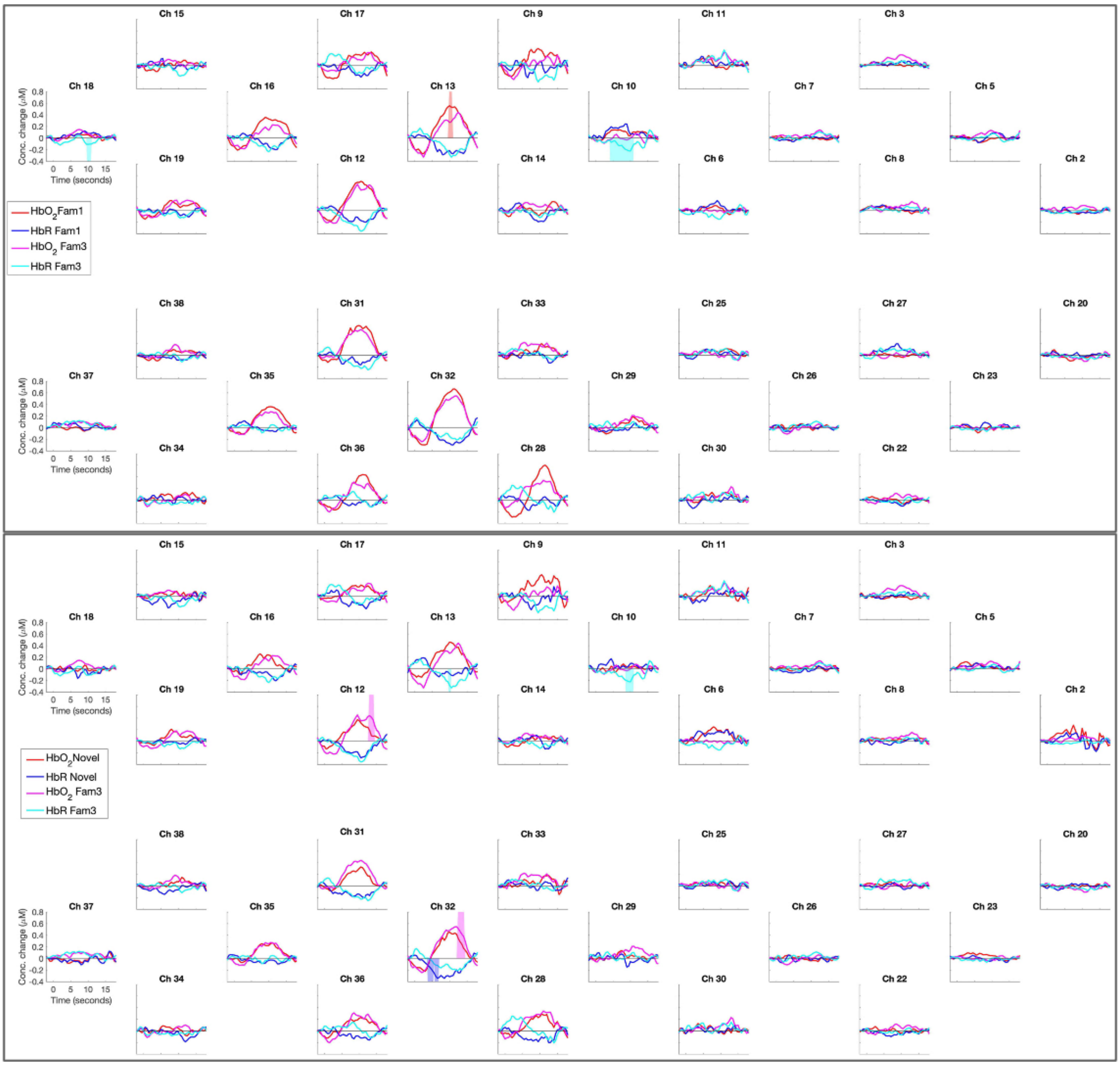


Figure S4. GM, 8mo visit: time courses of Fam1, Fam3 and Novel epochs. Upper panel: Fam1 vs Fam3 contrast. Lower panel: Novel vs Fm3 contrast. Shaded areas indicate significant difference between contrasts, the colour indicates which epoch is stronger (i.e. in the lower panel, magenta represents Fam3 HbO, then the magenta shaded areas represent Fam3 response is stronger than Novel).

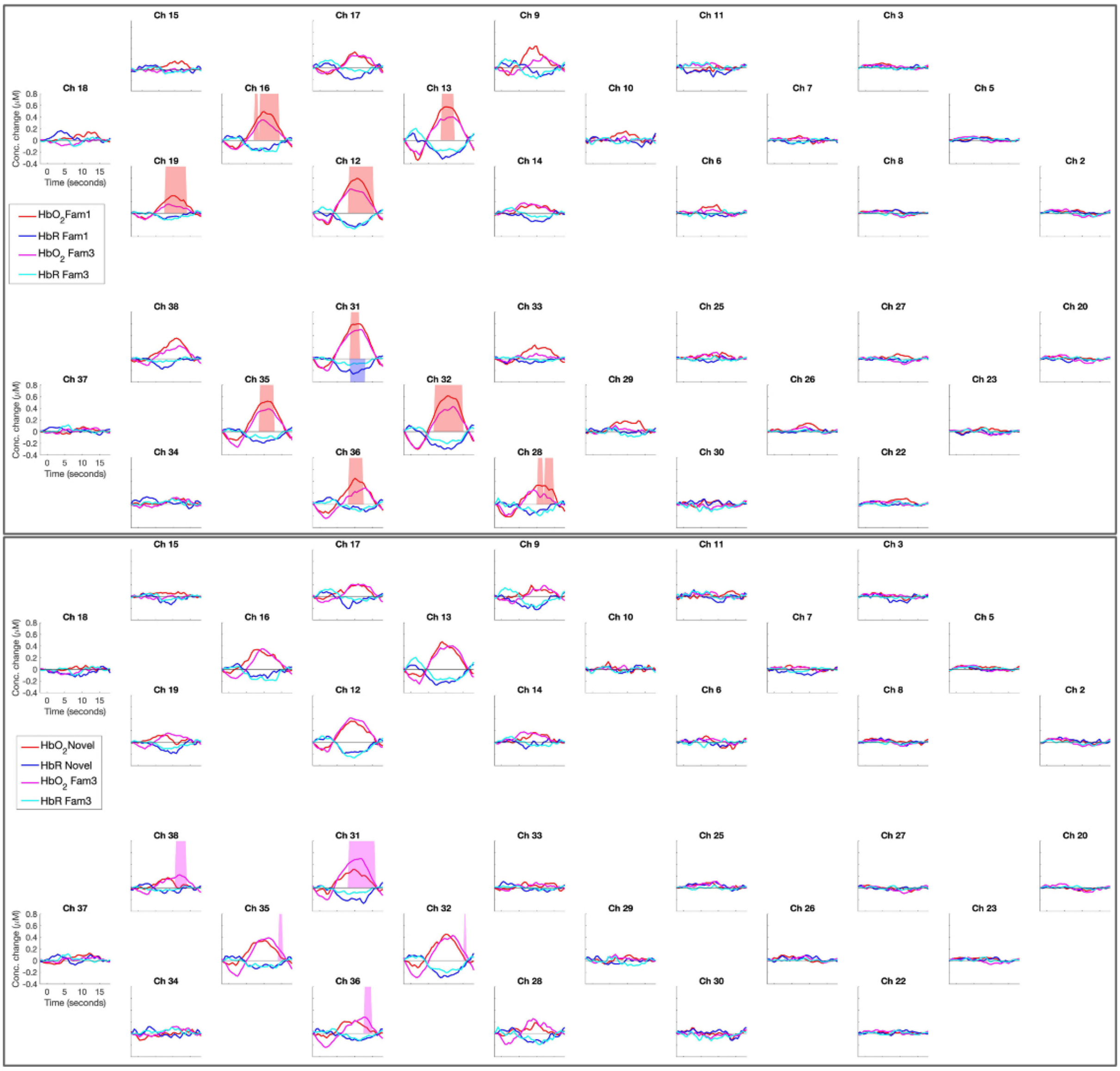


Figure S5. GM, 12mo visit: time courses of Fam1, Fam3 and Novel epochs. Upper panel: Fam1 vs Fam3 contrast. Lower panel: Novel vs Fm3 contrast. Shaded areas indicate significant difference between contrasts, the colour indicates which epoch is stronger (i.e. in the upper panel, red represents Fam1 HbO, then the red shaded areas represent Fam1 response is stronger than Fam3).

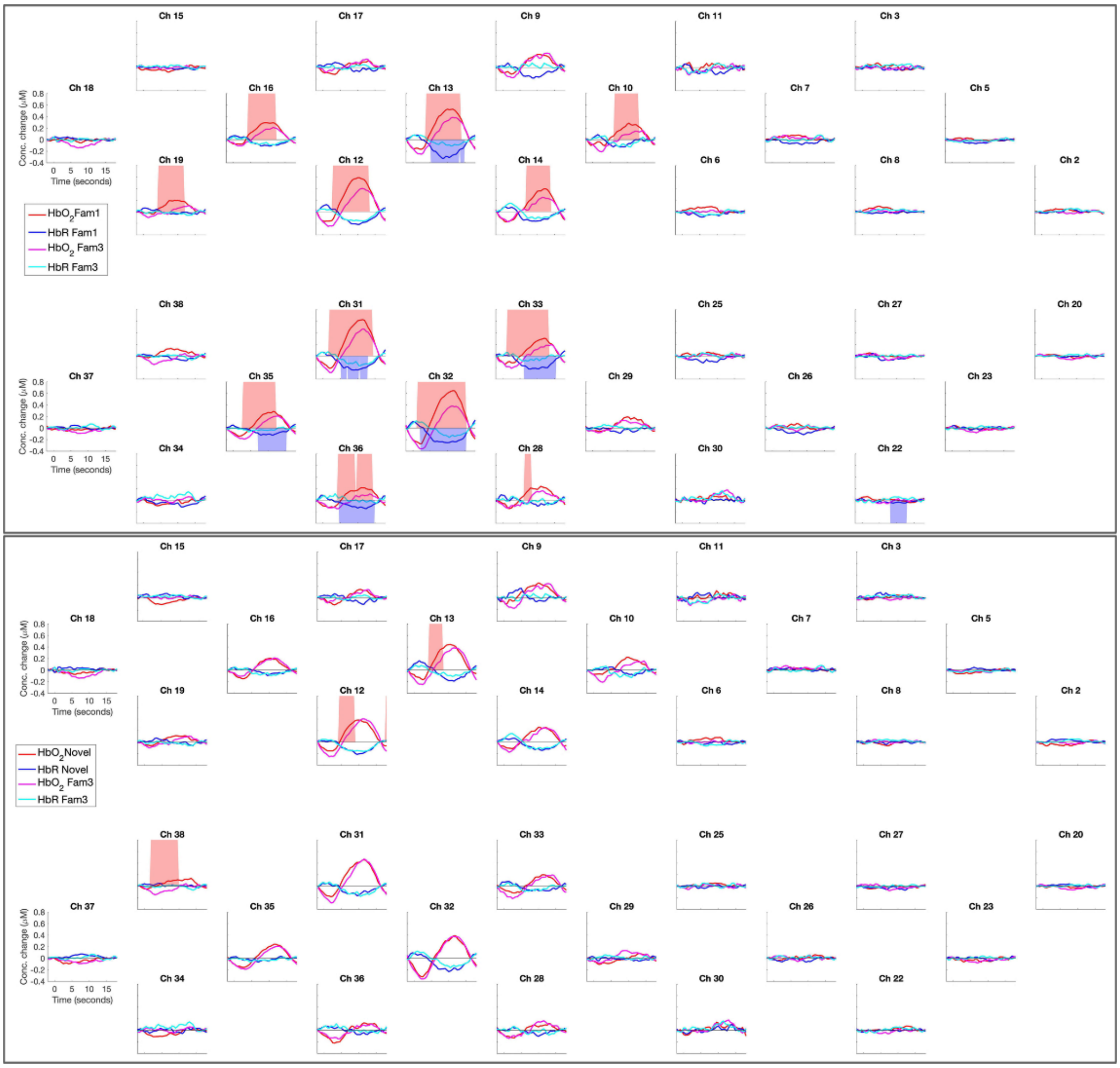


Figure S6. GM, 18mo visit: time courses of Fam1, Fam3 and Novel epochs. Upper panel: Fam1 vs Fam3 contrast. Lower panel: Novel vs Fm3 contrast. Shaded areas indicate significant difference between contrasts, the colour indicates which epoch is stronger (i.e. in the upper panel, red represents Fam1 HbO, then the red shaded areas represent Fam1 response is stronger than Fam3).

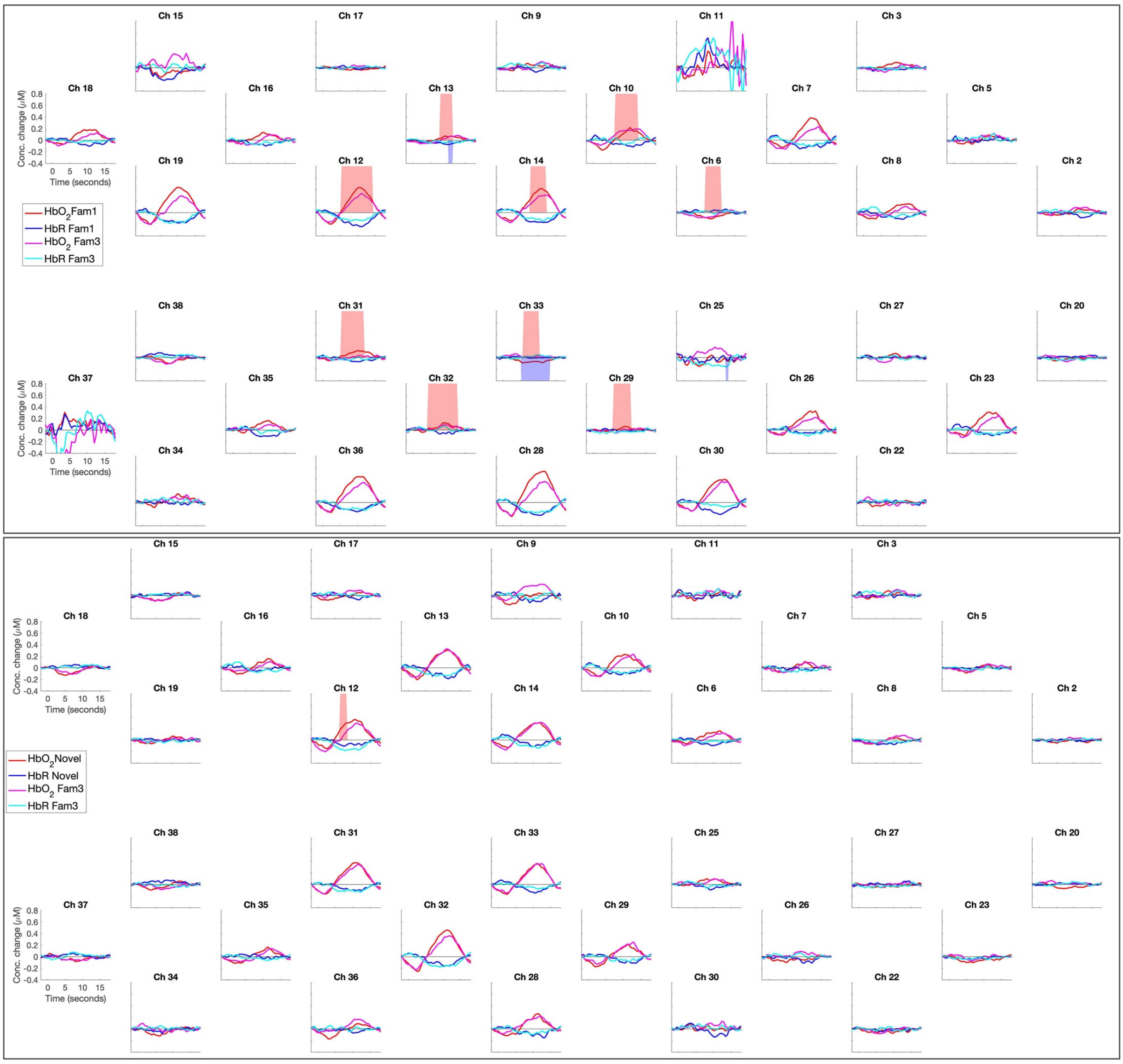


Figure S7. GM, 24mo visit: time courses of Fam1, Fam3 and Novel epochs. Upper panel: Fam1 vs Fam3 contrast. Lower panel: Novel vs Fm3 contrast. Shaded areas indicate significant difference between contrasts, the colour indicates which epoch is stronger (i.e. in the upper panel, red represents Fam1 HbO, then the red shaded areas represent Fam1 response is stronger than Fam3).

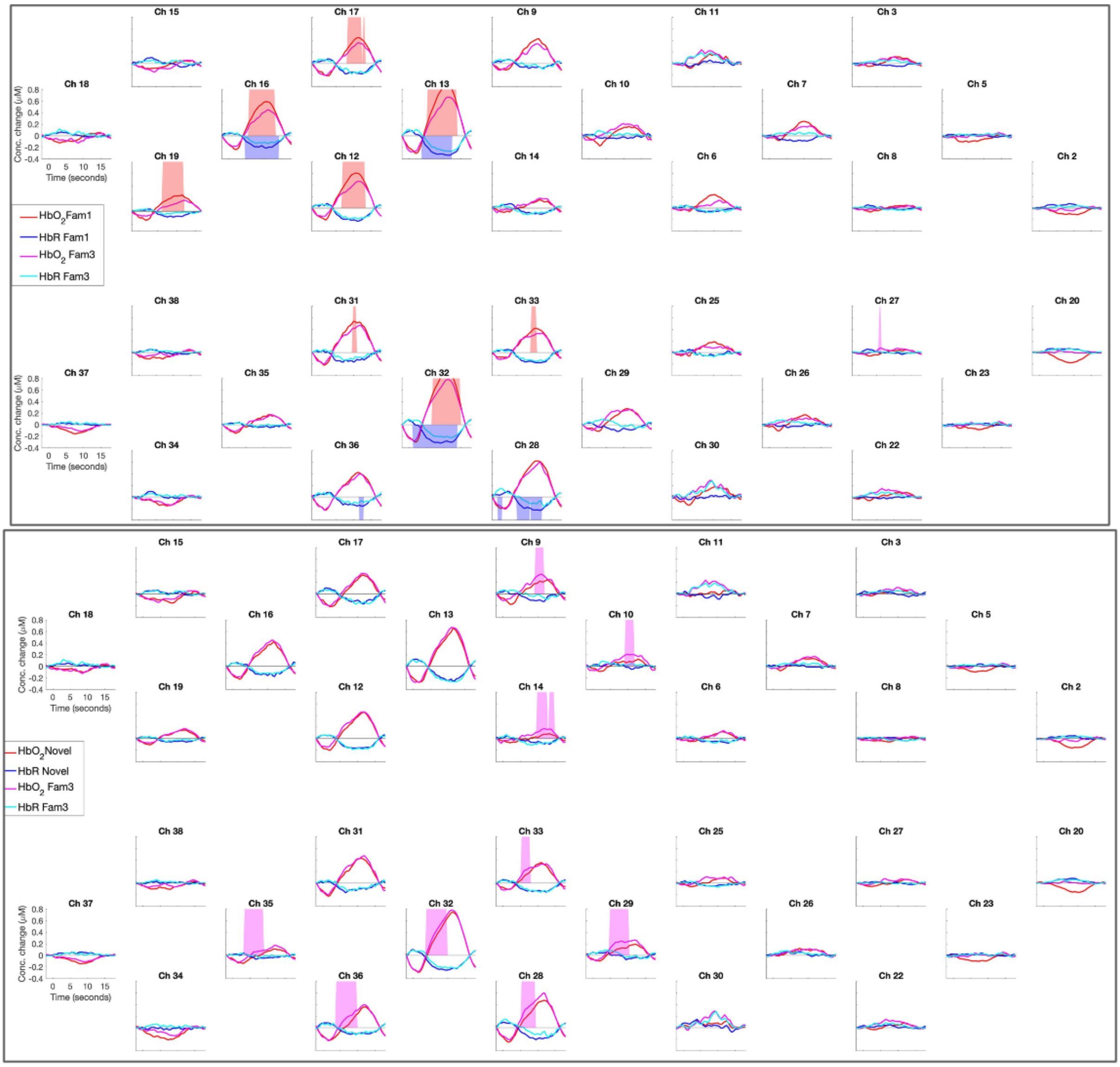


Figure S8. GM, 60mo visit: time courses of Fam1, Fam3 and Novel epochs. Upper panel: Fam1 vs Fam3 contrast. Lower panel: Novel vs Fm3 contrast. Shaded areas indicate significant difference between contrasts, the colour indicates which epoch is stronger (i.e. in the upper panel, red represents Fam1 HbO, then the red shaded areas represent Fam1 response is stronger than Fam3).

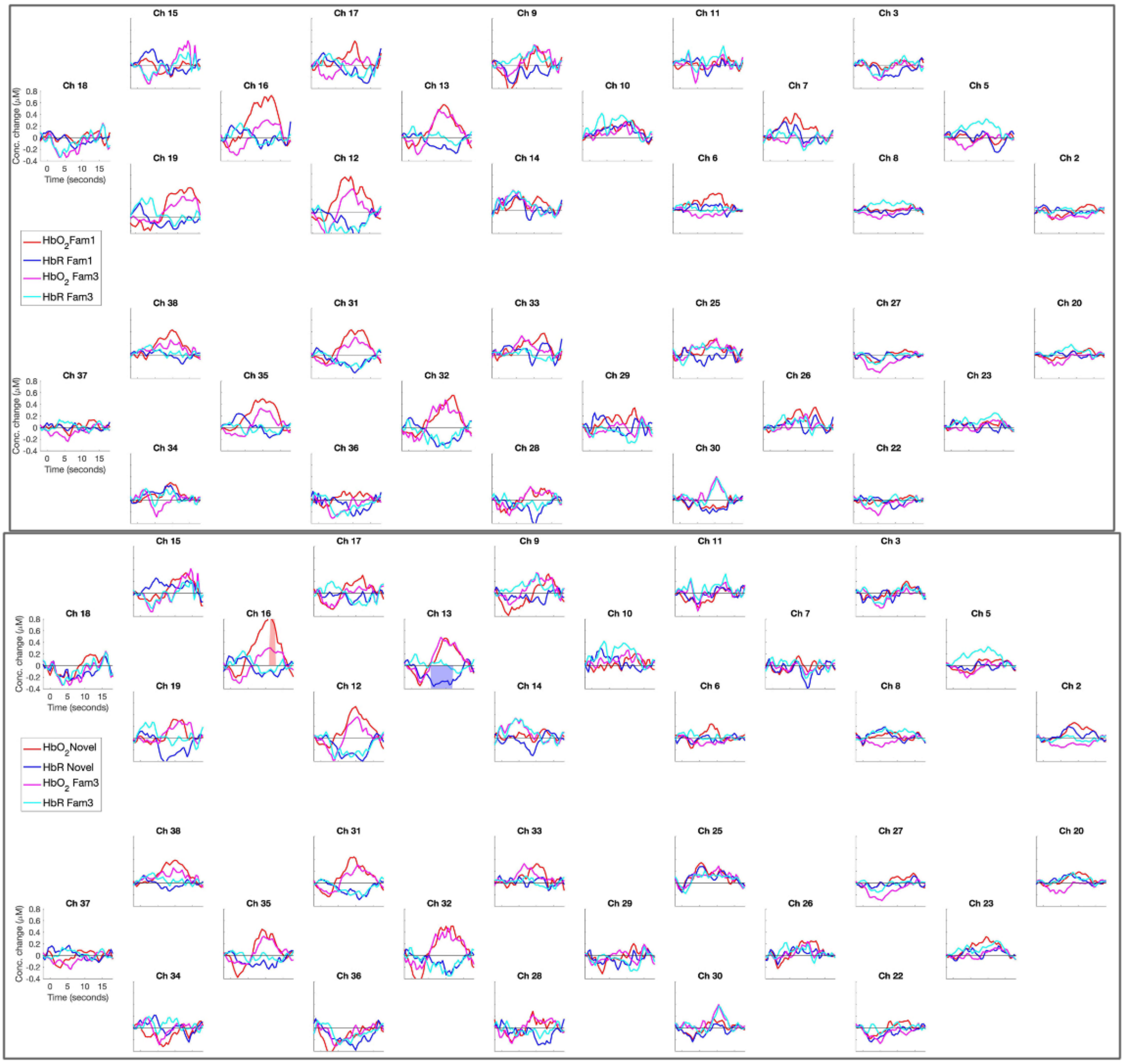


Figure S9. UK, 5mo visit: time courses of Fam1, Fam3 and Novel epochs. Upper panel: Fam1 vs Fam3 contrast. Lower panel: Novel vs Fm3 contrast.

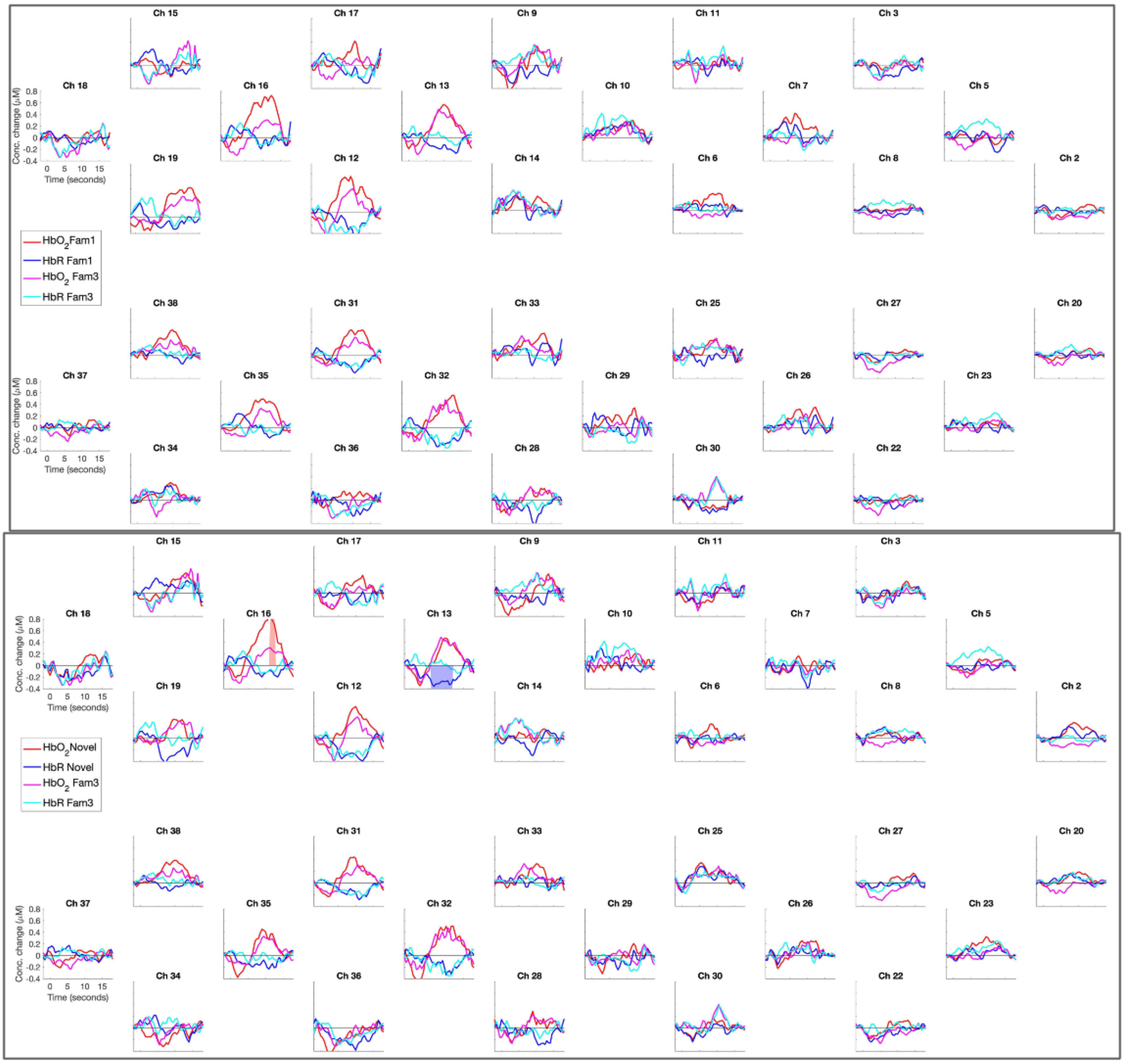


Figure S10. UK, 8mo visit: time courses of Fam1, Fam3 and Novel epochs. Upper panel: Fam1 vs Fam3 contrast. Lower panel: Novel vs Fm3 contrast. Shaded areas indicate significant difference between contrasts, the colour indicates which epoch is stronger (i.e. in the lower panel, blue represents Novel HbO, then the blue shaded areas represent Novel response is stronger than Fam3).

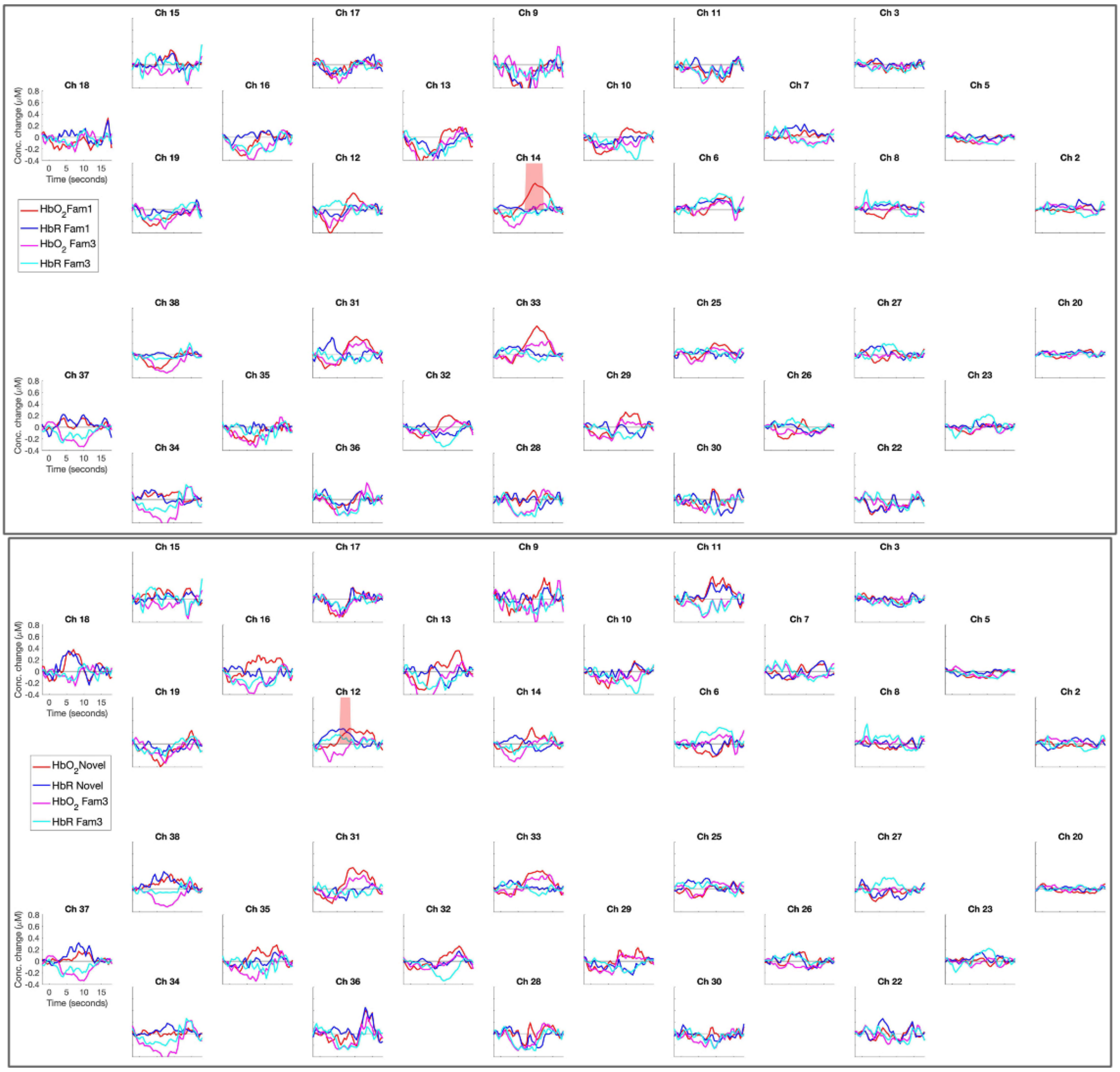


Figure S11. UK, 12mo visit: time courses of Fam1, Fam3 and Novel epochs. Upper panel: Fam1 vs Fam3 contrast. Lower panel: Novel vs Fm3 contrast. Shaded areas indicate significant difference between contrasts, the colour indicates which epoch is stronger (i.e. in the upper panel, red represents Fam1 HbO, then the red shaded areas represent Fam1 response is stronger than Fam3).

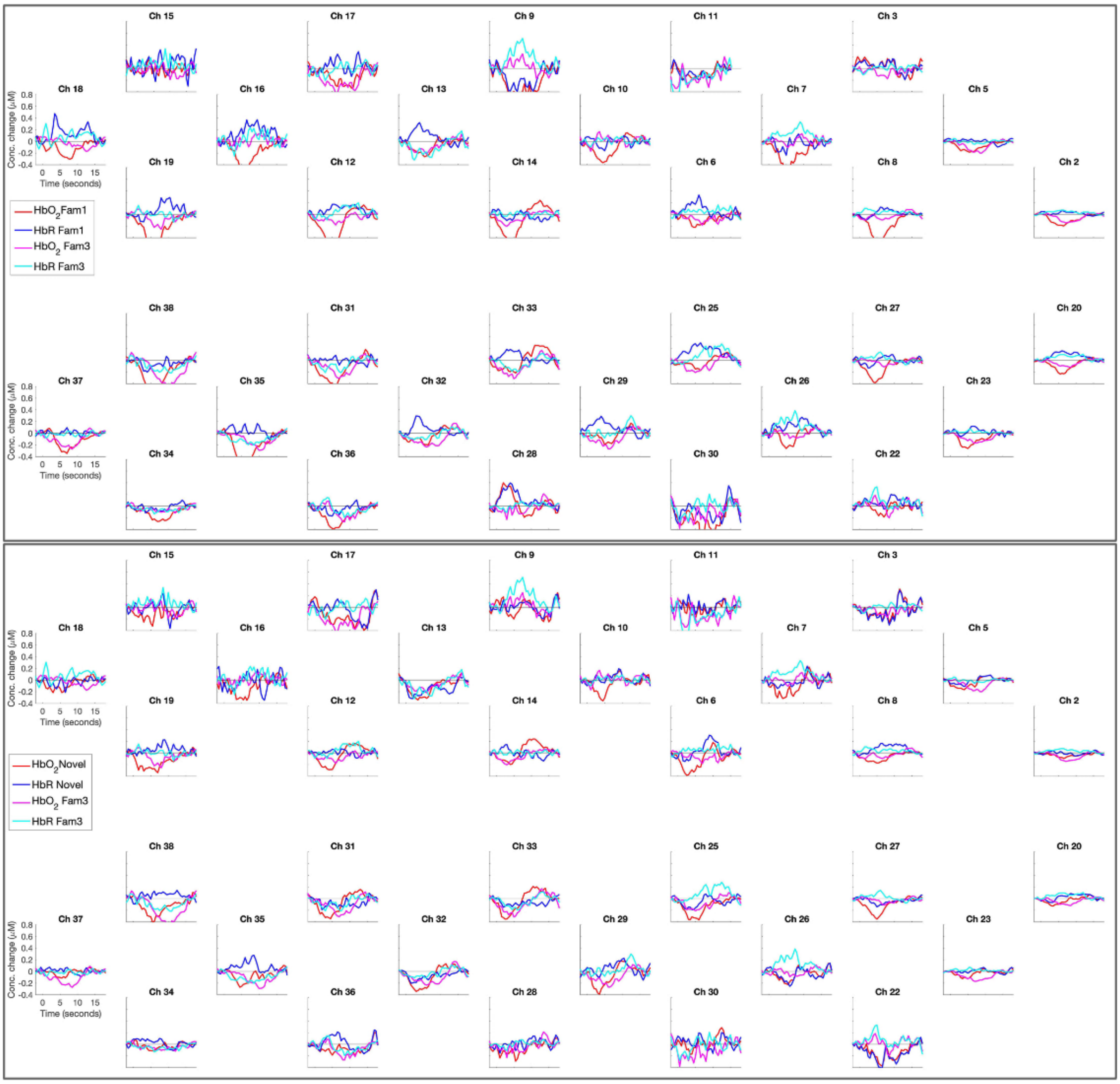


Figure S12. UK, 18mo visit: time courses of Fam1, Fam3 and Novel epochs. Upper panel: Fam1 vs Fam3 contrast. Lower panel: Novel vs Fm3 contrast.

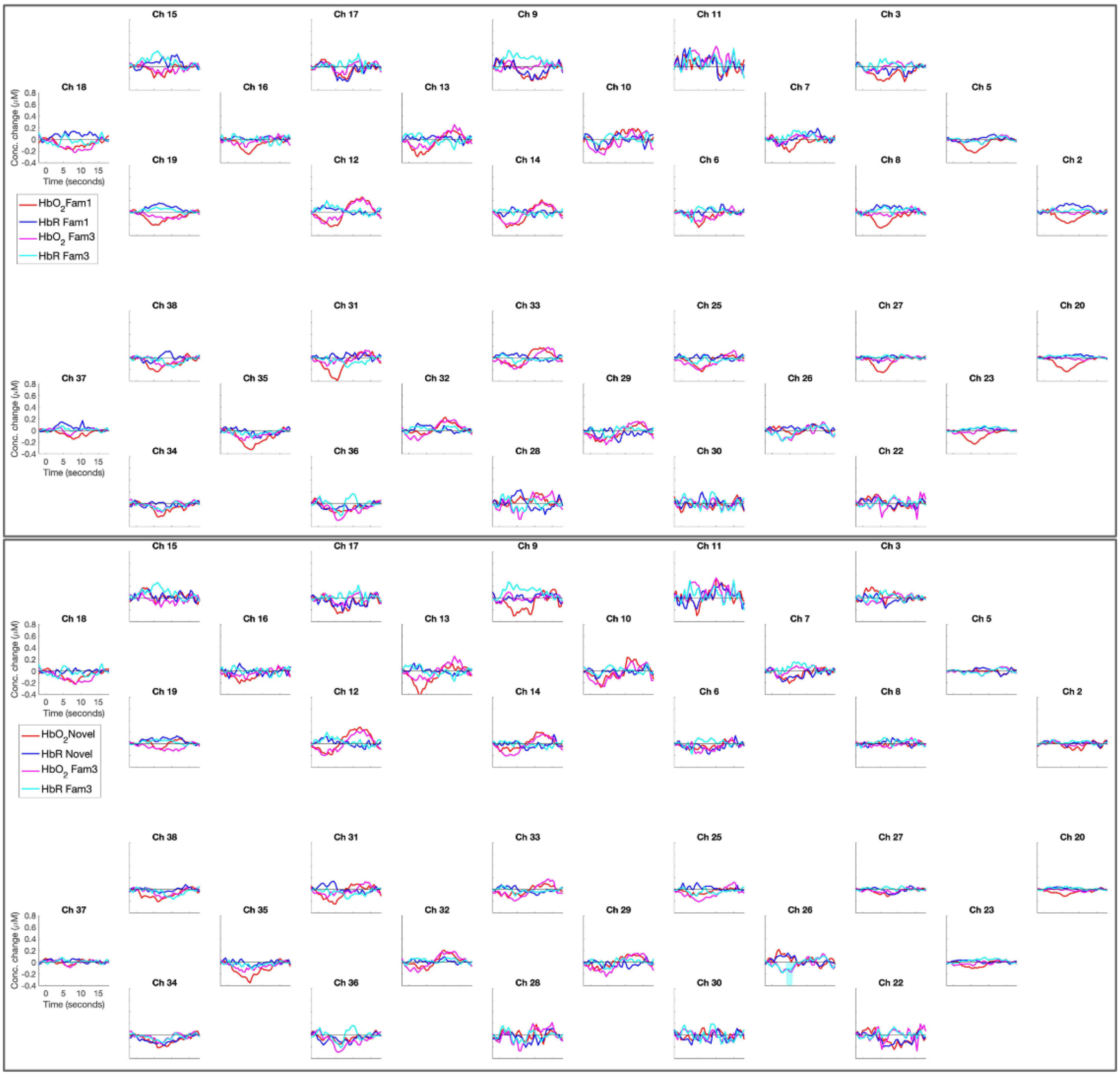


Figure S13. UK, 24mo visit: time courses of Fam1, Fam3 and Novel epochs. Upper panel: Fam1 vs Fam3 contrast. Lower panel: Novel vs Fm3 contrast.

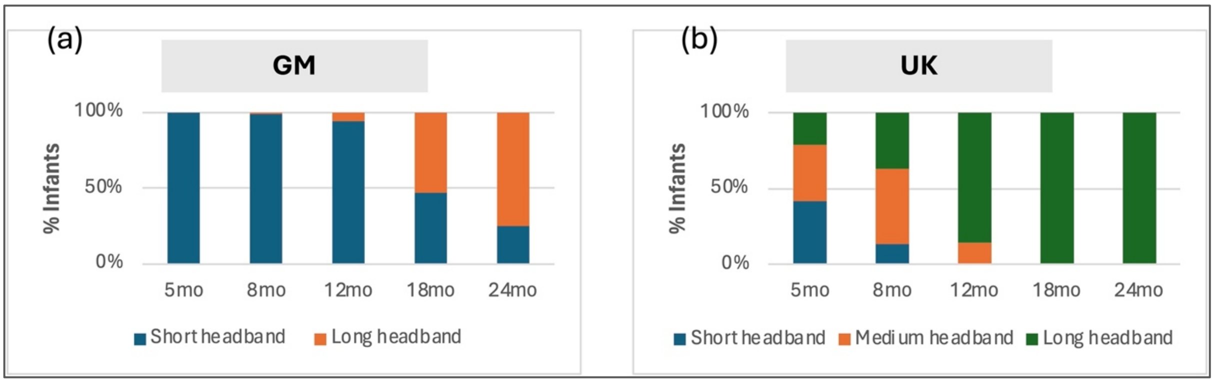


Figure S14. Percentage of participants using each size headband: (a) GM had two headband sizes: *short and long*; (b) UK had three headband sizes: *short*, *medium* and *long*.

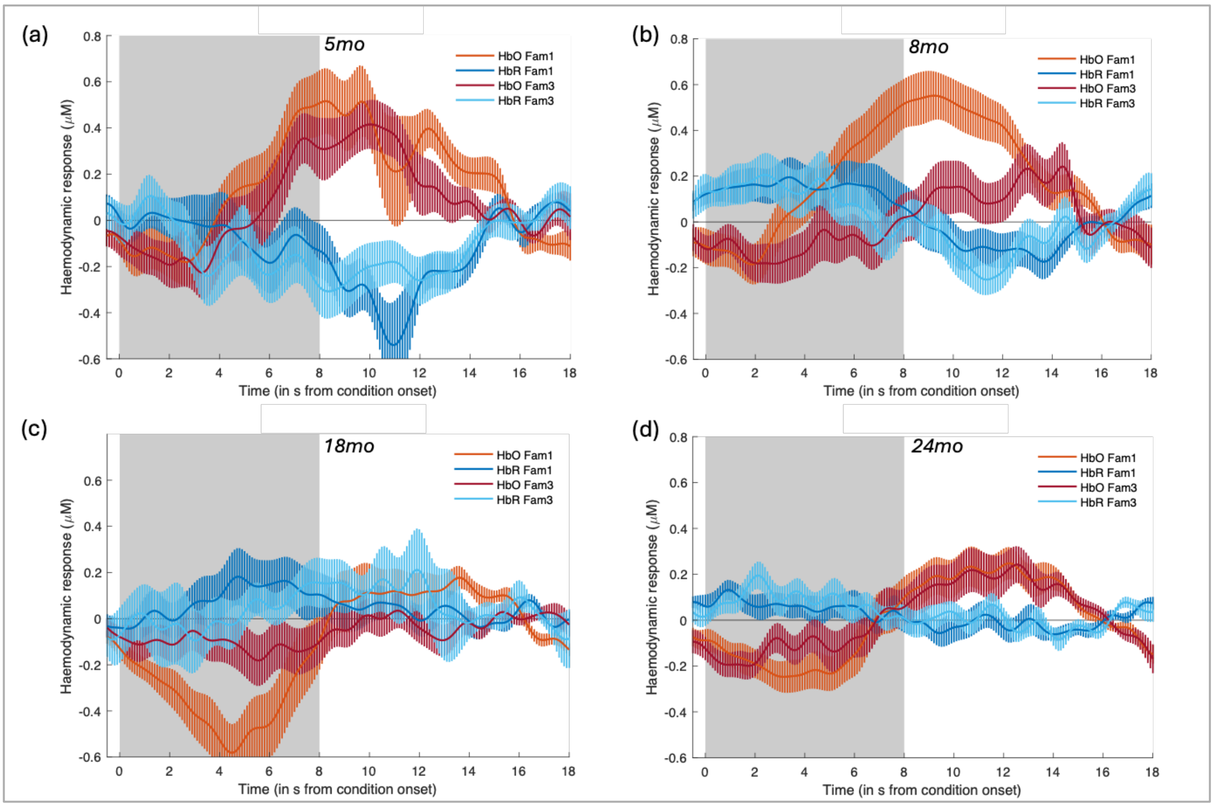


Figure S15. UK: Fam1 and Fam3 haemodynamic response in channel 12 at (a) 5mo, (b) 8mo, (c) 18mo and (d) 24mo. HbO2 time course (mean ± SEM) is represented in red and HbR in blue. The gray shaded area represents the condition presentation interval.
